# Complex multicellularity linked with expanded chemical arsenals in microbes

**DOI:** 10.1101/2025.08.06.668190

**Authors:** Rauf Salamzade, Lindsay R. Kalan, Cameron R. Currie

## Abstract

Single-celled microorganisms evolving to form cooperative multicellular units represents a key innovation in the history of life. Multicellularity is required for cell differentiation and, as such, is critical to the emergence of biological complexity. Here, we propose and find support for our hypothesis that multicellularity facilitated massive expansions in the biosynthetic potential to produce specialized metabolites across microbial life. Our systematic characterization of biosynthetic gene clusters (BGCs) across bacteria confirms that most taxa have limited predicted potential to produce natural products. In contrast with primarily unicellular lineages, we show that the origins of biosynthetic expansion in Actinomycetota, Cyanobacteriota, and Myxococcota coincide with independent origins of multicellular structures such as mycelia, filaments, and fruiting bodies. Further, we reveal that the two primary origins of expansions in specialized metabolism for fungi, corresponding to the Pezizomycotina and Agaricomycetes, similarly coincide with the emergence of complex multicellularity and fruiting body formation as well as mechanisms for self-identification during haplontic life cycles. Interestingly, all five of the biosynthetically talented multicellular microbial lineages are concomitantly enriched in carbohydrate utilization enzymes, suggesting that ancient chemical innovations have been further shaped by catabolic processes. Intraspecific cooperation shaping the evolution of specialized metabolism further informs our understanding of the major transitions in life and will help guide efforts to discover new antimicrobial compounds to counter the emergence of antibiotic resistance.

## INTRODUCTION

Evolving the ability to produce and secrete natural products, including antibiotics, represents a key innovation in microbial life^1,2^. Unlike primary metabolites essential for basic cellular functions, microbes leverage specialized metabolites to directly mediate complex interspecies interactions, compete for resources, and establish ecological niches^3^. Biosynthetic gene clusters (BGCs) encode the critical molecular machinery that enable microbes to produce the unparalleled chemical diversity present across microbially-derived natural products. By encoding complex enzymatic pathways for synthesizing bioactive compounds, BGCs have allowed microbes to evolve potent, diverse, and often highly specific metabolites that dramatically expand their functional capabilities to interact with and adapt to complex environmental challenges. They provide competitive advantages through various roles, including signaling^4^ and defense^2^. Microbial BGC-encoded natural products also represent a critical source of bioactive compounds with immense potential for drug discovery, having yielded numerous life-saving pharmaceuticals like antibiotics, immunosuppressants, and anticancer agents. Understanding the evolution of natural product synthesis by microbes is crucial for deciphering the role secreted metabolites play in mediating ecological interactions, shaping microbial diversity, and has transformative potential in drug discovery.

Multicellularity represents another major transition in the history of microbial life, enhancing biological complexity and enabling unprecedented diversification across multiple lineages^5^. The advantages conferred by multicellularity include improved resource acquisition, environmental resilience, and division of labor^6^. Multicellularity has emerged independently multiple times across the tree of life, including within the bacterial phyla Cyanobacteriota, Myxococcota, and Actinomycetota. Cyanobacteriota represent one of the earliest origins of multicellularity, with some species forming filamentous chains and differentiated cells that carry out specialized functions like nitrogen fixation in heterocysts^6^. The emergence of multicellularity in cyanobacteria has also been linked with taxonomic diversification and the Great Oxidation Event^7^. Some members of the bacterial phylum Myxococcota exhibit a form of collective behavior where individual cells coordinate to form complex multicellular structures called fruiting bodies in response to starvation conditions, with species like *Myxococcus xanthus* demonstrating sophisticated cell-to-cell communication and cooperative movement through social motility^6,8^.

The Actinomycetota represent a particularly fascinating example of bacterial multicellularity, with some taxa forming complex filamentous structures through hyphal growth that enables them to colonize and penetrate solid substrates like soil particles and plant material^9^. The multicellular lifestyle, as particularly evident in the genus *Streptomyces,* supports sophisticated developmental programs regulated by complex signaling networks, including the formation of aerial hyphae and spores in response to environmental cues^10^. Although other bacterial taxa can also exhibit allorecognition^11^ or form structured biofilms^12^, these traits often pale in complexity to the multicellular structures and life cycles of Actinomycetota, Myxococcota, and Cyanobacteriota or have emerged more recently and, as such, did not shape the genomic content of the deeper ancestral lineage from which it evolved.

Historically, the filamentous bacteria within the phylum Actinomycetota and the fungi within the phylum Ascomycota have served as the two most important microbial sources of natural products, contributing many important drugs, such as penicillin, cyclosporine, statins, streptomycin, tetracycline and erythromycin^9^. Ascomycetes and filamentous Actinomycetota exhibit complex life cycles during which they can form mycelia and intricate multicellular structures with distinct cell types as well as generate spores to survive stressful conditions and aid geographic dispersal to colonize new niches^9,13^. More recently, species belonging to the bacterial phyla Cyanobacteriota and Myxococcota have emerged as valuable and rich sources of novel natural products with biomedical potential^14–16^. In addition to containing taxa enriched in their ability to produce natural products, these microbial lineages also contain taxa that form cooperative multicellular units^6^. The concomitant occurrence of multicellularity within microbial lineages known to be enriched in the production of natural products suggests that the emergence of multicellularity facilitated the expansion of genomes and carriage of biosynthetic machinery. This is an intriguing idea that until now has not been examined across the domains of microbial life.

In this study, we systematically investigate biosynthetic capacities across the bacterial domain and fungal kingdom to assess the potential role of multicellularity in the enrichment of BGCs within phyla that have historically served as key sources of natural products. We highlight that the majority of bacterial and archaeal taxonomic orders are on average very limited in the number of specialized metabolites they encode per genome. In stark contrast, orders within the phyla Cyanobacteriota, Myxococcota, and Actinomycetota are broadly composed of extant genera rich in machinery for specialized metabolite production. Within all three phyla, the ancestral enrichment in biosynthetic repertoire coincides with early, independent origins of multicellularity. For Actinomycetota, we show the emergence of multicellularity and biosynthetic trait expansions align with the origin of the class Actinomycetes, the most sourced reservoir of bacterial natural products. We next investigate biosynthetic potential across the fungal kingdom and highlight that most lineages are limited in canonical specialized metabolism in comparison to the Dikarya lineages that are known for their formation of complex multicellular structures such as fruiting bodies. Through comparative and association analyses, we further show that increased biosynthetic capacity across microbial lineages is strongly associated with enrichments in carbohydrate-active enzymes as well as proteins mediating self-recognition in fungi. Specialized metabolites and carbohydrate-active enzymes released into the extracellular environment constitute public goods that can generate tragedy of the commons dynamics within microbial populations. Multicellular organization reduces selection on individual cells to pursue their own self-interest, making cooperative production of these energetically costly molecules more evolutionary stable.

## RESULTS

### Ancestral expansions in biosynthetic capacity are limited across the bacterial tree of life

To begin evaluating the association of multicellularity with enrichments in specialized metabolism, we first determined distributions of per-genome biosynthetic carriage for 74 well-sequenced bacterial and archaeal orders. Both the median number of BGCs as well as the median BGC-ome size, the summed length of predicted BGC regions in a single genome, were assessed across 4,732 genus-representative genomes from the orders (**Fig. 1a; Supplementary Table S1**). A strong correlation was observed between the two metrics and while complete genomes were preferentially used where available, BGC-ome size was largely assessed throughout this study to alleviate potential over-estimation of biosynthetic capacity related to assembly fragmentation issues. Trends in BGC-ome size were also consistent when requiring stricter criteria for annotating BGCs (**Extended Data Fig. 1**). The distribution of biosynthetic genomic content was heavily skewed, with the vast majority (65 of 74) of bacterial and archaeal orders having a median count of five or fewer BGC regions or a maximum median BGC-ome size of less than 150 kb (**Fig. 1a**). None of the eight archaeal orders had a median count of five or more BGCs, confirming their known lack of biosynthetic traits^17^. Large-scale annotation efforts have also previously suggested that BGC carriage is limited in many bacterial species^18^ and databases cataloging the taxonomic origins of known natural products have shown that they are disproportionately sourced from just six microbial phyla^19,20^ (**Fig. 1b**).

**Fig. 1:**
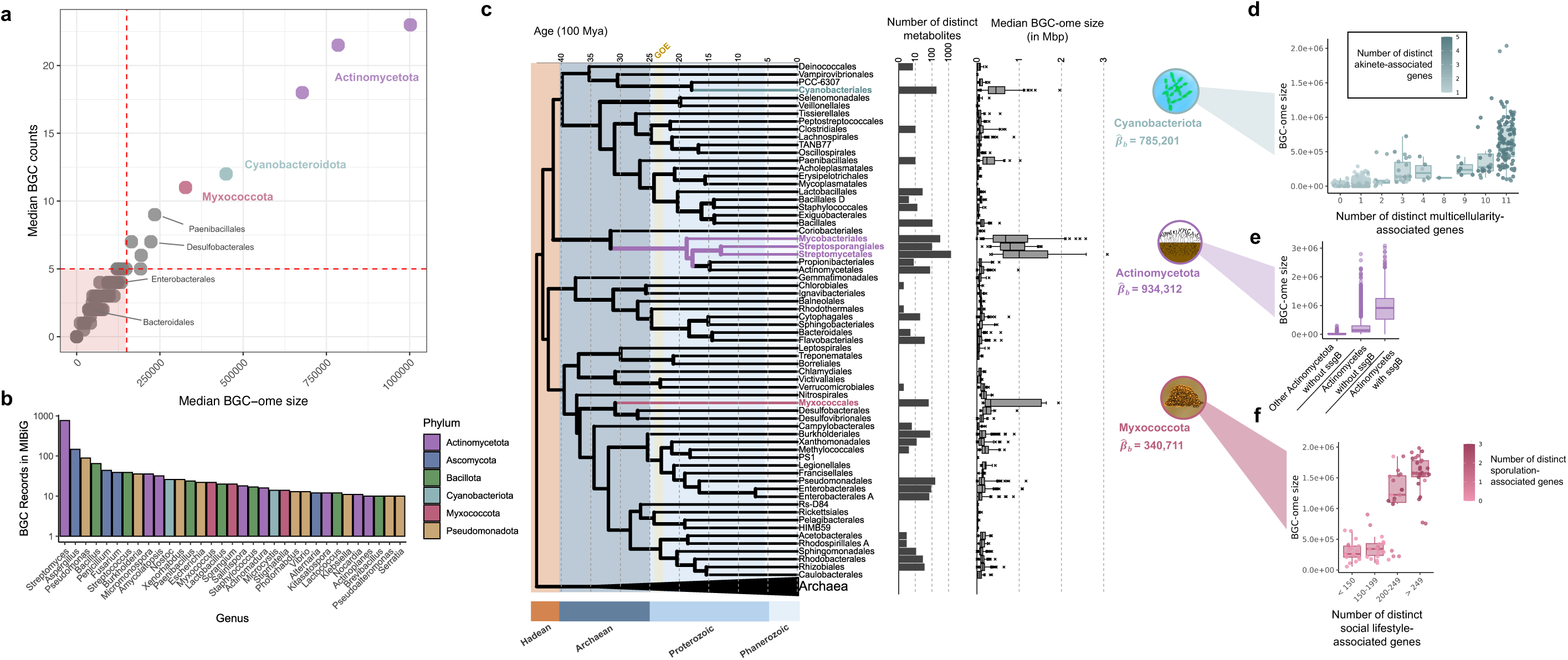
Ancestral expansions in biosynthetic content coincide with the developments of multicellularity. a, The median count of BGC regions and median BGC-ome size for 74 bacterial and archaeal orders. **b,** Counts of characterized BGCs for all bacterial or fungal genera in MIBiG v3.1 with >10 characterized BGC records. **c,** A timetree of 74 bacterial and archaeal orders. The bar chart depicts the number of distinct metabolite types (distinct nodes from NPAtlas) for each order. Box-plots depict the count of total BGCs for representative genomes from each order. The coloring for highlighted orders coincide with panel **a.** The width of the branches corresponds to the standard deviation of time estimates, with greater variability shown as bolder branches. Evolutionary expansions in median BGC-ome size for each order detected using the Ornstein-Uhlenbeck process are shown with the magnitude of the shift 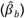 indicated to the right. **d**, The relationship between the number of distinct multicellularity genes from Boden et al. 2023 to BGC-ome size for species-representative Cyanobacteriota genomes. The number of akinete related genes described by Singh, Khan, and Srivastava 2020 is shown as the coloring. **e**, The BGC-ome sizes for Actinomycetes with and without SsgB, a genetic marker for mycelium formation, alongside BGC-ome sizes for other classes from Actinomycetota. **f**, The relationship between the number of distinct social-lifestyle related genes from Murphy et al. 2021 to BGC-ome size for species-representative Myxococcota genomes. The number of sporulation specific proteins identified by Dahl et al. 2007 is shown as the coloring.

To establish which taxonomic orders exhibit substantial increases in their median BGC-ome size relative to their phylogenetic contexts, we next constructed a time tree of the 74 bacterial and archaeal orders and applied a phylogeny-informed approach leveraging Ornstein-Uhlenbeck processes (**Fig. 1c; Supplementary Table S2**). This analysis highlighted five distinct orders belonging to three phyla: Cyanobacteriota, Myxococcota and Actinomycetota. Because each of the three phyla are enriched in biosynthetic content across multiple genera, we posit that they correspond to cases of ancestral enrichments in biosynthetic traits. As described in the introduction, each phylum also features some of the most well-characterized model bacterial species used for studying complex multicellularity and life strategies. Four other bacterial orders have median counts of six or more BGCs per genome and also include species that exhibit multicellularity-like or cooperative behavior. They include the orders of Desulfobacterales, which feature multicellular magnetotactic bacteria^21^ and Paenibacillales, which encompasses species that have complex physiologies and can perform organized colony swarming^22^.

### Ancestral enrichments in bacterial biosynthetic capacity are associated with multicellularity, sporulation, and carbohydrate active enzymes

We next investigated whether a variety of genomic, functional and ecological traits correlate with increased biosynthetic capacity across the three phyla highlighted as having undergone ancestral biosynthetic expansions, in addition to the phyla of Pseudomonadota and Bacillota (**Supplementary Notes; Extended Data Fig. 1-2; Supplementary Table S3**). Across 17,844 species-level representative genomes from the phyla, multiple genomics inferred features were positively correlated with BGC-ome size, including: (i) carriage of oxidative phosphorylation-related enzymes, (ii) increased coding usage of specific amino acids and (iii) an inferred preference for mesophilic growth. After genome size, enrichment for carbohydrate processing enzymes was most strongly associated with increased BGC-ome size, including in Pseudomonadota and Bacillota. Further, multiple independent genome-wide association studies (GWAS) across phyla revealed that several plant-related carbohydrate usage enzyme families were significantly associated with biosynthetic capacity. The only family found to be associated with BGC-ome size in four or more of the five phyla was glycosyl hydrolase family 19, chitinases that are rare in microbes and hypothesized to have been horizontally acquired in *Streptomyces* from plants^23^. Thus, these analyses support a significant increase in biosynthetic capacity broadly occurred across diverse bacteria due to interactions with and usage of biopolymers from land plants^24^.

Within the three phyla determined to have had ancestral expansions in biosynthetic content, positive associations were observed between counts for known protein markers of multicellularity and sporulation^25–27^ with BGC-ome size (**Fig. 1d-f**). For Cyanobacteriota, carriage of eight or more multicellularity-associated genes and three or more akinete-associated genes were significantly associated with increased BGC-ome size (*p*=2.59e-59). Similarly, for Myxococcota, genomes with greater than 200 social-lifestyle-associated genes and two or more sporulation-associated genes had larger BGC-ome sizes (*p*=4.65e-13). The substantial overlap in social lifestyle-associated orthologs among genomes spanning diverse taxonomic orders further suggests sociality was ancestral to the phylum Myxococcota or, at least, multiple classes within it (**Extended Data Fig. 3).** Amongst Actinomycetota, the carriage of *ssgB* homologs, which are involved in the process of sporulation and associated with increased genome size and complex physiology^26^, was also a clear indicator of increased biosynthetic potential (*p*<1e-300). Further, many species within the class Actinomycetes, even those without *ssgB* homologs, had larger BGC-omes in comparison to other classes in the phylum (*p*<4.50e-126; **Fig. 1e**).

### The last common ancestor of Actinomycetes was multicellular and possessed a large reservoir of BGCs

To further understand the enrichment of biosynthetic content and emergence of multicellularity in the class Actinomycetes we first used phylogenomics to determine the evolutionary relationship between 133 genera with high-quality genomes from across the phylum Actinomycetota (**Fig. 2a; Supplementary Table S4**). A maximum-likelihood phylogeny was constructed based on single-copy-core ortholog groups revealing two clades within the Actinomycetes class, we refer to as clade-1 and clade-2 (**Fig. 2ab; Supplementary Table S4**). Based on literature^9,28^, both clade-1 and clade-2 were found to feature prominent filamentous and mycelium-forming genera and also found to genomically encode proteins associated with multicellularity, such as SsgB, SepX, RamA and RamB^26,29,30^.

**Fig. 2:**
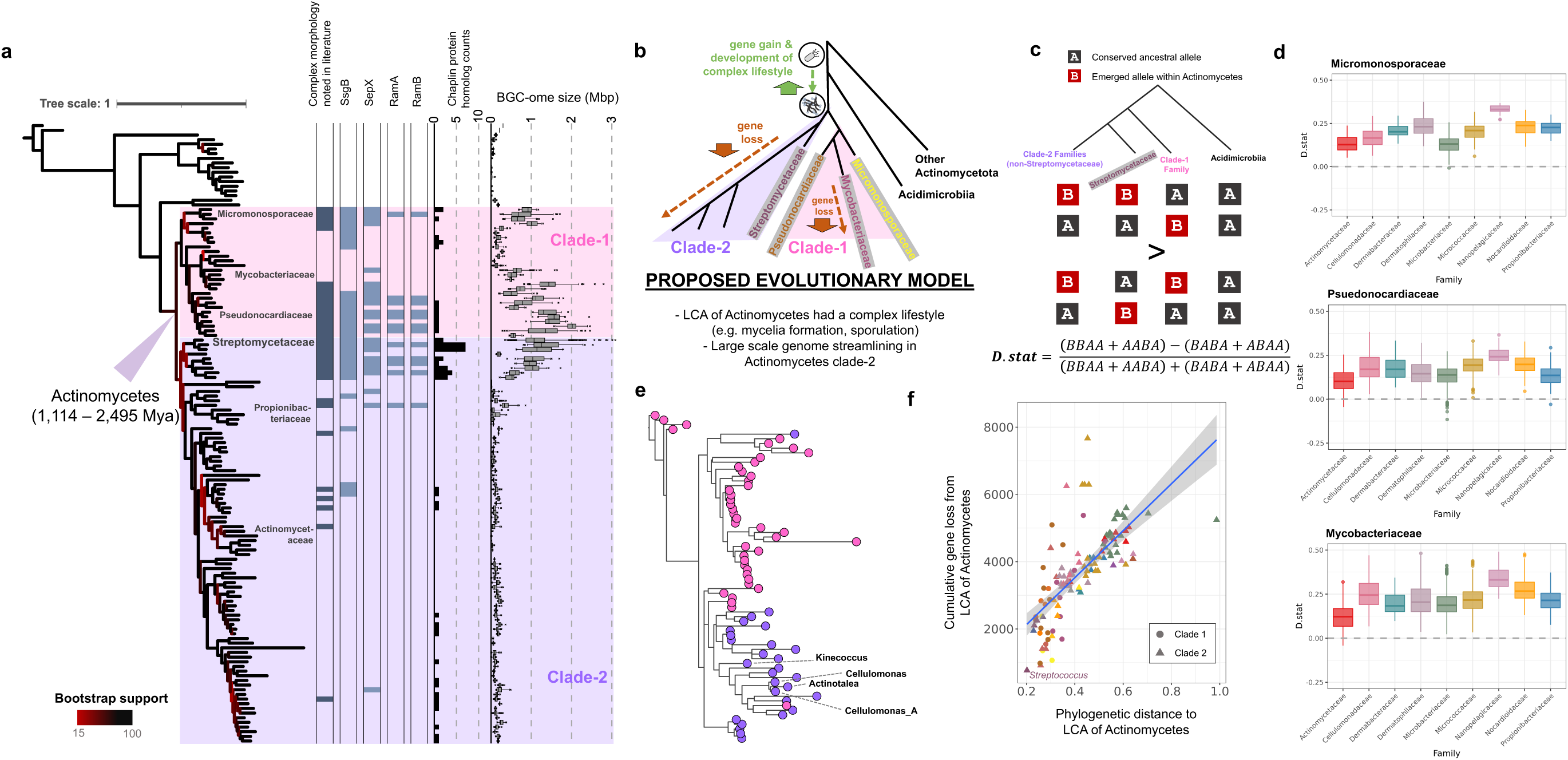
The last common ancestor of Actinomycetes was morphologically complex and possessed a rich biosynthetic repertoire. **a**, A maximum-likelihood phylogeny of representative genera from the Actinomycetota phylum from conserved single-copy ortholog groups. The Actinomycetes class is highlighted in violet and pink, indicating two major ancestral partitions observed for the class. The presence of ortholog groups corresponding to genetic markers associated with multicellularity, sporulation, or mycelium formation in Actinomycetota, primarily in *Streptomyces*, are shown across the phylogeny of the full phylum. The set of boxplots furthest to the right showcases the BGC-ome size for species representative genomes from each genus. The estimated age interval for the LCA of Actinomycetes correspond to the 89% high density interval predicted using MCMCTree. **b,** A schematic of the proposed evolutionary model of Actinomycetota, **c,** Overview of the D.stat for inferring whether smaller genome Actinomycetes in clade-2 share a more recent common ancestor with Streptomycetaceae than with families belonging to clade-1. **d,** Distributions of the D.stat computed for the single-copy core alignment with different combinations of families from clade-2 (boxplots) and clade-1 (grids). **e,** Maximum phylogeny of SsgB proteins with sequences from smaller genomes (< 6 Mbp) from clade-2 marked. **f,** Phylogenetic distance of Actinomycetes genomes to the last common ancestor node in the phylogeny correlates with estimated cumulative gene loss from ancestral state reconstruction inference.

The phylogenetic positioning of filamentous genera suggests that multicellularity was ancestral to the Actinomycetes class. This is supported by multicellularity being present across most clade-1 lineages as well as clade-2 featuring the early-diverging and filamentous family of Streptomycetaceae. This suggests that morphologically simpler genera from clade-2, like *Micrococcus*, have experienced greater gene loss and a loss of physiological complexity. While ancestral loss of morphological complexity has previously been suggested for the order of Mycobacteriales in clade-1^10^, to our knowledge, it has not been hypothesized to have occurred at large-scale, beyond one order^31^, within clade-2 Actinomycetes. Due to low bootstrap values at certain ancestral nodes in our phylogenomic model, we further assessed the proposed topology for Actinomycetes evolution using several additional phylogenomic and evolutionary approaches (**Supplementary Notes; Fig. 2c-f; Extended Data Fig. 4**). This process included developing a statistical framework to assess highly conserved sites along conserved and single-copy orthologs, which revealed consensus support that unicellular taxa from clade-2 share a more recent last common ancestor with Streptomycetaceae than with morphologically complex families from clade-1.

A multicellular last common ancestor of Actinomycetes coinciding with the broad enrichment of biosynthetic traits in its extant genera further supports that a large biosynthetic repertoire was ancestral to this class. Genome streamlining in less morphologically complex taxa across Actinomycetes likely explains why BGCs are elevated across the class but natural products discovery has still been largely limited to filamentous actinomycetes^20^ (**Extended Data** Fig. 5a). Some BGCs were also likely lost in these genera together with their ability to filament and form mycelia. To assess the conservation of specialized metabolism across the phylum Actinomycetota, we assessed ortholog groups found to be key components of BGCs in two or more Actinomycetes. Of the 428 biosynthetic ortholog groups identified, the majority, 366 (85.5%), were found within BGC contexts in both major clades of the class (**Fig. 2g; Extended Data Fig. 5b; Supplementary Table S5**). Further, of those 366 ortholog groups, 266 (72.7%) were either exclusive to Actinomycetes or exclusively found within a BGC context within Actinomycetes. This includes the ortholog group corresponding to 3-dehydroquinate (DHQ) and aminoDHQ synthases, involved in shikimate and 3-amino-5-hydroxybenzoic acid (AHBA) biosynthesis, respectively. Phylogenetic investigations further revealed that one key enzyme in AHBA biosynthesis, aminoDAHP synthase, emerged within Actinomycetes and evolved from type II 3-deoxy-7-phosphoheptulonate (DAHP) synthases involved in the shikimate pathway (**Supplementary Notes; Expanded Data Fig. 6**). AHBA is a key precursor of the functionally vast ansamycin class of specialized metabolites^32^, which the antibiotic rifamycin belongs to.

While rifamycin resistance is widespread across bacteria^33^, we found that protein markers associated with it amongst Actinomycetota were largely specific to Actinomycetes and common across clade-1 and clade-2 of the class, including many genera lacking morphological complexity (**Supplementary Table S6**). We next performed a targeted search for homologous instances of rifamycin-synthesizing BGCs across the entire phylum. All such instances were from Actinomycetes and exhibited high sequence conservation, despite belonging to diverse genera spanning both clades of the class and support that their evolution has largely occurred vertically (**Supplementary Notes; Expanded Data Fig. 6**). Together these results suggest ansamycin and AHBA biosynthesis machinery were encoded by the last common ancestor of Actinomycetes and further demonstrate how biosynthetic gene clusters can arise from genes involved in primary metabolism^34^.

### Significant enrichments in biosynthetic traits for fungi coincide with developments in complex multicellularity and haplontic life cycles

A hallmark of fungal natural products is the Ascomycota, the source of critically important drugs such as the antibiotic penicillin, the immunosuppressant cyclosporine and the cholesterol-lowering statins. Like the other major microbial source of natural product drugs, the Actinomycetota, the Ascomycota contain genera enriched in BGCs and have the ability to form complex cooperative units in the form of mycelia, suggesting that enrichment in biosynthetic potential in fungi is associated with multicellularity. To systematically investigate BGC enrichment across fungi, we first constructed a high-resolution phylogeny of the kingdom using highly conserved genes to assess the relationship of 312 genomes, including several recently sequenced early-diverging taxa^35^ (**Fig. 3; Extended Data Fig. 7a; Supplementary Table S7-S8**).

**Fig. 3:**
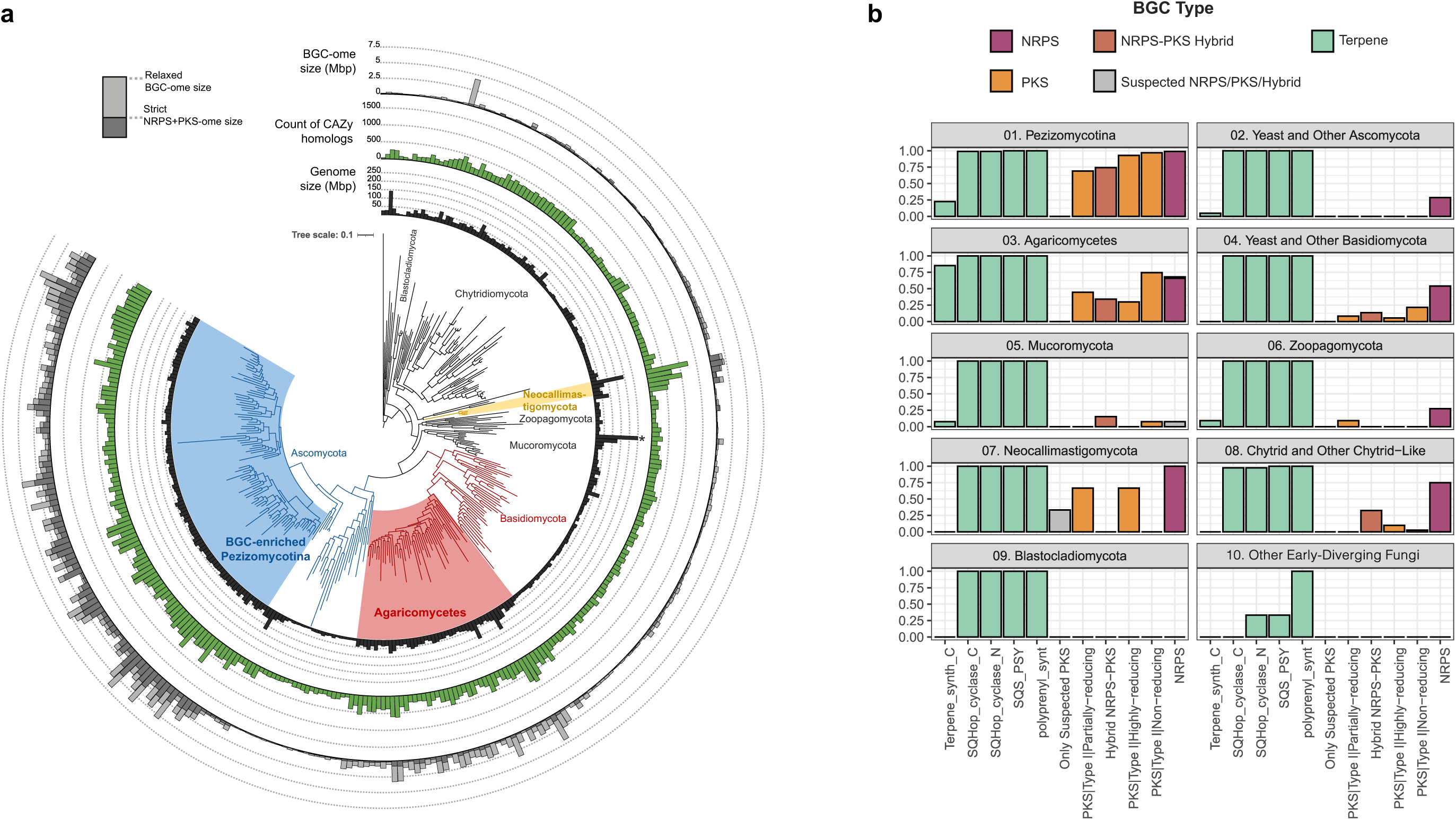
Phylogenetic overview for enrichments in BGC-ome size across the kingdom of fungi. a, A maximum-likelihood phylogeny of fungi rooted using outgroup genomes based on marker genes from the UFCG database. The innermost barplot (black) shows the genome size for representative genomes. Note, the genome size for *Gigaspora rosea* (522.9 Mbp) was capped at 250 Mbp for visualization. The middle barplot in green shows the number of total CAZy enzyme homologs found across genomes and the final barplot (in shades of grey) shows the proportion of genomes predicted to be BGCs. The BGC enriched clades for Chytridiomycota and related phyla, Ascomycota, and the Basidiomycota are highlighted in gold, red, and blue, respectively. **b,** The occurrence of NRPS and PKS types along with certain terpene synthesis related domains is shown across different fungal taxonomic clades. The category ”Other Early-Diverging Fungi” orrespond to *Rozella allomycis*, *Paramicrosporidium saccamoebae*, and *Mitosporidium daphniae*.

Next, we conducted comparative genomics exploring the biosynthetic repertoire across the fungal tree of life. As with bacteria, we show that most fungal lineages are composed of taxa with limited biosynthetic potential, with ∼77% of distinct non-dikaryotic genera having ten or fewer BGC regions or BGC-ome sizes of 500 kb or shorter (**Fig. 3a**). Among fungi, two lineages (*Rozella*, *Paramicrosporidium*, and *Mitosporidium*) and the Blastocladiomycota, have the smallest BGC-ome sizes and lack polyketide synthases (PKS) and non-ribosomal peptide synthetases (NRPS) (**Fig. 3b**). These trends and the lack of BGCs in *Fonticula*, a close outgroup of the fungi, support the conclusion that limited biosynthetic potential is symplesiomorphic in fungi. Fungal genomes from the Chytridiomycota, Mucoromycota, and Zoopagomycota all have relatively small BGC-ome sizes but contain a diverse set of terpene domains as well as some PKS and NRPS BGCs, including those substantially diverged from sequences underlying the synthesis of known specialized metabolites (**Fig. 3b; Extended Data Fig. 7bc**). Interestingly, the phylum of Neocallimastigomycota exhibited enrichment of BGCs relative to chytrids and other non-dikaryon fungi (*p*=2.48e-3; **Extended Data Fig. 7d**). These anaerobic fungi commonly inhabit the guts of herbivores and were recently noted for their high biosynthetic potential due to gene acquisition from bacteria via lateral transfer events^36^. They can also form morphologically complex rhizoids^37^.

Next, in our evaluation of the evolution of biosynthesis in fungi, we find support for two independent origins of BGC enrichment within the Dikarya, each coinciding with the emergence of complex multicellularity from morphologically less-complex yeasts. Specifically, phylogenetic analyses indicated that minimal BGC enrichment is symplesiomorphic in the Dikarya, with the sister lineage to Dikarya, the Mucoromycota, and early-diverging lineages of both the Ascomycota and Basidiomycota having relatively small BGC-ome sizes. Amongst Ascomycota, a monophyletic subdivision within the Pezizomycotina was determined to be greatly enriched in BGCs relative to all other fungi (*p*=2.34e-29; **Fig. 3a**; **Extended Data Fig. 7e**), a result previously identified by others^38,39^. While hyphal multicellularity evolved early in fungal evolution^40^, consistent with our observations in bacteria, the ancestral expansion of biosynthetic content in the Pezizomycotina coincides with developments in complex multicellularity, such as the formation of fruiting bodies. Within Basidiomycota, the class Agaricomycetes represents a second origin of BGC enrichment with taxa generally featuring larger BGC-ome sizes than other Basidiomycota species as well as non-dikaryotic fungal species (*p*=3.87e-12 & *p*=6.68e-14, respectively; **Fig. 3a; Extended Data Fig. 7f**). As with the Pezizomycotina, the Agaricomycetes represent the other major fungal group well known for possessing complex multicellularity^41^, including the ability to produce dikaryons^35,42^.

We next investigated which orthologous families of proteins associate with increased biosynthetic capacity in fungi. To do so, we performed comparative genomic and GWAS (**Fig. 4a**). One trait revealed as enriched in BGC-rich dikaryotic fungal clades are proteins implicated in functioning as heterokaryon incompatibility (HI) factors. Heterokaryon or vegetative incompatibility (HI) proteins enable self-identification amongst filamentous fungi^43,44^ and mechanistically involve autophagy, which are key properties of many multicellular organisms. A sensitive search for domains associated with HI^45^ revealed that many species within BGC-enriched Pezizomycotina and Agaricomycetes feature hundreds of proteins with these domains in their genomes (**Fig. 4bc, Extended Data Fig. 8a-e**). Interestingly, biosynthetic regions of BGC-rich Pezizomycotina had slight, but significant, elevated rates of HI proteins (*p*=6.59e-4), suggesting a possible connection between BGCs and vegetative incompatibility in these fungi. The correlation between BGC-ome size and HI counts was further supported at the genus-resolution within *Aspergillus* (**Extended Data Fig. 8f-k; Supplementary Table S11**). Outside of these two clades, other fungi typically dedicated a significantly lower proportion of their total proteins to HI (*p*=1.40e-37; **Extended Data Fig. 8e**).

**Fig. 4:**
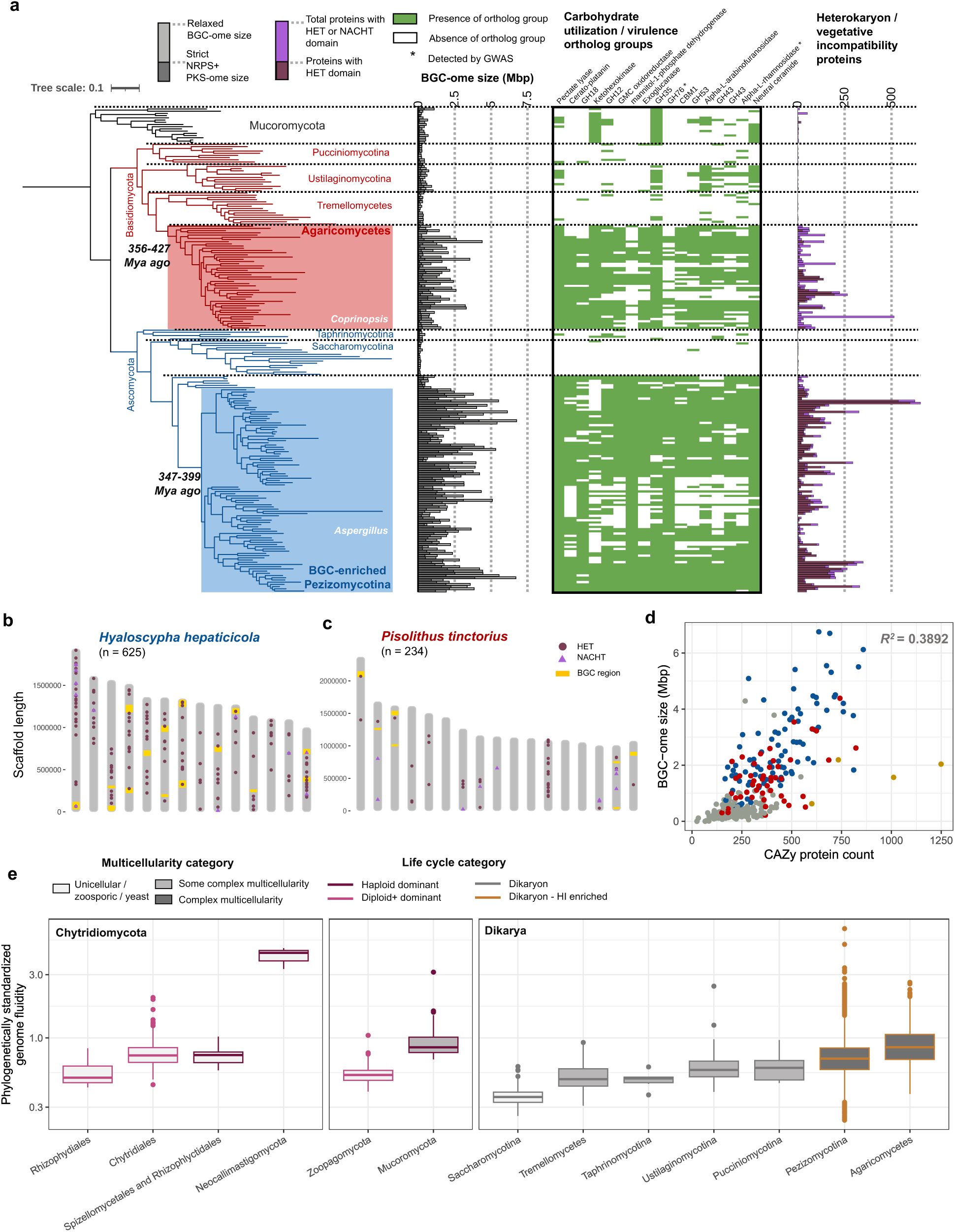
Associated genetic factors with biosynthetic potential across fungi. a, The phylogenetic distribution of select ortholog groups associated with carbohydrate utilization or virulence as well as the number of proteins annotated as featuring domains related to heterokaryon / vegetative incompatibility proteins are shown across Dikarya. The estimated age intervals for the LCA of the BGC-enriched Pezizomycotina and Agaricomyetes correspond to the 89% high density interval predicted using MCMCTree. **b,** Boxplots of pairwise genome fluidity values standardized for phylogenetic distance between genomes. Values were capped at 10.0. Coloring of boxplot frames corresponds to life cycle category of the taxonomic groups and the fill indicates whether they are known to exhibit complex multicellularity. The distribution of BGCs and HI-associated proteins across scaffolds ≥ 1 Mb in **c,** *Hyaloscypha hepaticiola* and **d,** *Pisolithus tinctorius.* Values indicated in parentheses correspond to the total number of suspected HI-associated proteins for each genome. **e,** The association of carbohydrate utilization protein counts is shown with BGC-ome size. Genomes are colored in accordance with clade coloring in **Fig. 3a**. R2 values were computed using linear regressions adjusted for phylogenetic relationships between genomes.

Given their function, the enrichment of HI proteins across Pezizomycotina supports that they typically exist in a vegetative state as haplontic homokaryons^35,42^. To assess the genomic impacts of life cycles, we next developed a phylogenetically standardized measure of genome fluidity and used it to determine that clades with greater genomic fluidity tended to be haplontic (**Fig. 4e; Extended Data Fig. 9a**). Similar trends were observed for BGC-ome size distributions, with haplontic clades generally exhibiting greater biosynthetic potential (**Extended Data Fig. 9b**). Despite having fewer BGCs, Agaricomycetes might simply appear to have more expansive pangenomes than Pezizomycotina because they are more likely to be in a dikaryotic form and harbor two distinct genomes at the time of sampling for genome sequencing^35^.

Our investigations further revealed that several specific ortholog groups predicted to function in carbohydrate processing or plant-virulence are either associated with increased biosynthetic capacity across fungi or enriched in BGC-rich Pezizomycotina and Agaricomycetes relative to other Dikarya (**Fig 4a; Supplementary Table S9-S10**). More broadly, carbohydrate processing enzyme enrichment was more strongly associated with biosynthetic capacity than even genome size in fungi (**Fig. 4d; Extended Data Fig. 7gh**). Thus, provided that both Pezizomycotina and Agaricomycetes likely originated around the same time as the emergence of land plants^46^, intricate relationships with plants are likely associated with biosynthetic enrichments in the two dikaryotic clades in addition to developments in multicellularity and mechanisms for self-recognition.

## DISCUSSION

The production of bioactive specialized metabolites is a hallmark of microbial diversity and drug discovery. Here, we systematically confirm that most lineages of microbial life carry a limited number of BGCs, as inferred by genomic annotation of canonical biosynthetic content. We then show that in three bacterial phyla and two fungal classes signatures of ancestral expansions in biosynthetic traits coincide with early developments of multicellularity. Further, our robust phylogenomic analyses infer that the last common ancestor of the class Actinomycetes, formerly Actinomycetia, which features many biosynthetically gifted genera including *Streptomyces,* was multicellular and potentially able to form aerial hyphae. While hyphal formation is likely ancestral to the fungal subkingdom Dikaryota^40^, complex multicellularity, such as fruiting body formation, likely arose multiple times in the ancestors of the discrete, biosynthetically rich, sublineages Pezizomycotina and Agaricomycetes. The broad co-occurrence of complex multicellularity with BGC enrichment in extant lineages of these bacterial and fungal taxa suggests intraspecific cooperation helped spur ancestral expansions in biosynthetic capacity to produce specialized metabolites, including natural products that serve as key drugs in human health.

The benefits microbes derive from producing natural products are obvious. These potent bioactive molecules mediate complex intra- and inter-species interactions within free-living and host-associated microbial communities, including in the human microbiome. However, these specialized metabolites require complex enzymatic machinery and are energetically expensive compounds to produce. Multicellular organization likely helps mitigate the fitness costs associated with production of specialized metabolites through division of labor and facilitates the evolution of expanded genomes encoding more BGCs. Likewise, filamentous growth would facilitate an individual microbe reaching the biomass required to produce natural products at ecologically relevant concentrations. The higher biomass obtained from multicellular growth would also facilitate the use of secreted carbohydrate-active enzymes to deconstruct the more complex biopolymers that would otherwise be unavailable, such as chitin and cellulose, which would concomitantly support the biosynthesis of bioactive molecules to suppress competitors for these energy rich substrates. Indeed, our finding that BGC expansions in both fungi and bacteria coincides with significant enrichment in carbohydrate utilization enzymes suggest that catabolic processes help these microbes obtain the energy required to produce specialized metabolites at ecologically relevant scales. Furthermore, microbes and their specialized metabolites have been shown to participate in intricate relationships with terrestrial and marine eukaryotic hosts, such as defensive symbiosis^47,48^. Taken together, our work supports a model in which several biosynthetic expansions have occurred over geological time, with an earlier expansion coinciding with early developments in multicellularity and a more recent expansion owing to emergent relationships with algae, invertebrates, and plants. The subsequent expanded access to energy through the use of carbohydrate active enzymes likely further fueled developments in more complex multicellular structures, offset costs of carriage of a greater arsenal of specialized metabolites, and promoted the transition from diffusion-feeding life histories to invasive growth strategies.

In addition to carbohydrate utilization machinery, we highlight the high prevalence of proteins suspected to function in HI in the two BGC-rich fungal clades of Pezizomycotina and Agaricomycotina. HI proteins allow filamentous dikaryons to self-recognize and initiate autophagy to prevent the formation of heterokaryons. The ability to self-distinguish is likely particularly important for fungi being able to outcompete other fungal species for access to potentially limited carbon sources within their natural habitats. The high copy counts of Hi proteins in Pezizomycotina is perhaps expected considering that the dominant life stage of these filamentous ascomycetes is as haplontic homokaryons. However, it is surprising that suspected HI proteins, in particular NLR-related ones, are so prevalent in Agaricomycetes which commonly exist as dikaryons in a sexual life cycle^35^. Recently, NRL-related HI proteins in fungi were shown to share ancestry with antiphage proteins from bacteria, which were found in higher copy counts for multicellular phyla, including Actinomycetota, Myxococcota, and Cyanobacteriota^49^. It is thus interesting to posit that this family of proteins might also serve more fundamental roles in enabling multicellularity by supporting cellular differentiation or compartmentalization in both bacteria and fungi, perhaps as a foundational mechanism for autophagy. Furthermore, since haplontic Pezizomycotina are more biosynthetically gifted than Agaricomycetes, both within Dikarya and broadly across fungi, haplontic life cycles are associated with increased biosynthetic capacity and perhaps also more expansive pangenomes. The basis for these trends remains to be further studied with one possible reason being that meiotic recombinations during non-haplontic life cycles breaking up linkage between epistatic alleles, including within BGCs.

Specialized metabolites must be secreted to mediate interactions within an organism’s ecological niche, and as such, operate as ’public goods’^50^. The production of costly public goods makes cells vulnerable to exploitation by ‘cheater’ cells that benefit from the specialized metabolite without investing metabolic resources in their production, setting up ‘tragedy of the commons’ dynamics. Some unicellular bacteria employ quorum sensing systems to help resolve tragedy of the commons dynamics; these chemical communication networks enable synchronized group behaviors only when sufficient population density is reached, thereby preventing premature and inefficient production of costly public goods. Given that in multicellular organisms all cells are genetically identical or clonally related, the high relatedness reduces the incentive to ‘cheat’ because benefits of cooperation flow back to genetically identical cells. As such, the resulting kin selections conferred by multicellularity avoids the tragedy of the common dynamics known to cause breakdowns in cooperative behavior as individual cells pursue their own interest, rather than producing and secreting natural products.

The evolution of multicellularity presents a fundamental paradox in evolutionary biology, as it requires individual cells to sacrifice their direct reproductive fitness to benefit the collective organism. Our findings suggest that a major benefit conferred by the emergence of multicellularity is concomitant ability to evolve a high capacity to produce potent chemical mediators of intra- and interspecific interactions. Further, these chemical compounds may serve in yet unrecognized roles in mediating within-species dynamics, further enforcing cooperative dynamics within multicellular entities, as hinted at by our finding of abundances of HI-associated proteins within BGC-rich and fruiting body-forming fungi. Our findings provide new insights into the evolution of multicellularity and suggest that multicellular taxa, especially those that are understudied and/or from less commonly investigated habitats, might be prime sources for the discovery of truly novel classes of specialized metabolites.

## ACKNOWLEDGEMENTS

This work was supported by grants from the National Institutes of Health [NIAID U19AI142720 (C.R.C & L.R.K) and NIGMS R35GM137828 (L.R.K.)]. C.R.C. received partial support from the Stephen A. Jarislowsky Foundation and the Canadian Institute for Advanced Research. The content is solely the responsibility of the authors and does not necessarily represent the official views of the National Institutes of Health. The authors would like to thank Timothy James, John Taylor, Ted Shultz, Adam Schaenzer, Gerard Wright, Asaf Salamov, Marguerite Langwig, Grant Nickles, and Soleil Young for helpful discussions, feedback or advice.

## AUHTOR CONTRIBUTIONS

C.R.C. conceptualized the origin of the study. R.S. curated the data; performed formal analysis; developed genomic methodologies. R.S., L.R.K., and C.R.C. designed the study; wrote the original draft of the article; developed visualizations; and reviewed and approved of the final manuscript.

## COMPETING INTERESTS

The authors declare no competing interests.

## METHODS

### Code availability

Programs, scripts, commands, configuration files, and select datasets for the study can be found in the GitHub repository: https://github.com/Kalan-Lab/Ancestral_BGC_Exansions_Study which is also version tracked and stored on Zenodo [add Zenodo citation]. The codoff program used in part to infer whether BGCs are from bacterial contamination in fungal genomes is available at: https://github.com/Kalan-Lab/codoff. The paiof (& hamtree) programs, used to profile the amino acid identity distributions between paired genomes and construct neighbor-joining trees based on ortholog group carriage information following OrthoFinder analysis, are provided in the GitHub repository: https://github.com/raufs/paiof. The psaps program to assess pangenome expansion rate differences between fungal clades is available on the GitHub repository: https://github.com/raufs/psaps. Releases of all GitHub repositories are also tracked in immutable forms on Zenodo.

### Data availability

NCBI accession codes for genomic assemblies used for different analyses presented in this study are provided in **Supplementary Table S1, S3, S4, S6, S7, and S11**. Larger datasets, including select files from antiSMASH BGC predictions, and fungal gene calling data can be found on the Zenodo repository: https://zenodo.org/uploads/15168076.

### Construction of a bacterial and archaeal timetree and determination of evolutionary shifts in BGC counts

A bacterial and archaeal timetree was constructed for 74 representative bacterial and archaeal orders which have been well-sequenced. To construct the phylogeny a two-tiered approach was used. First, high-resolution phylogenies were constructed for each of the five bacterial phyla with the most number of characterized BGCs based on predetermined gene sets shown to be largely single-copy and highly conserved for bacteria^51,52^ using IQ-TREE (v2.2.0.3)^52^. These were assessed to determine more reliable topologies for individual phyla than what could be achieved with a more limited number of single-copy core genes that are more broadly conserved across both bacteria and archaea. To construct the final phylogeny of the 74 representative bacterial and archaeal orders, a guiding tree constraining intra-phylum topologies and specifying a partition between Terrabacteria and Gracilicutes^53^ was provided to IQ-TREE (v1.6.12) together with trimmed protein alignments of eight conserved ribosomal proteins^54^ which featured a combined total of 1,095 amino acid sites and were conserved and non-ambiguous in ≥90% of genomes. IQ-TREE was run requesting an edge-unlinked partition model with ModelFinder limited to the LG and WAG substitution matrices.

Next, to establish a timetree, we followed the protocol described by Dos Reis 2022 with some adaptations to parameter configurations^55^. Fossil and prior-inference based calibration information for node ages were gathered from literature^56,57^ (**Supplementary Table S2**). These calibrations were encoded as upper and lower bounds for select nodes in the phylogeny provided as input for MCMCtree (v4.10.0)^58^. antiSMASH (v7.0.0)^59^ was used for BGC annotation of genus-level representative genomes. Finally, to determine evolutionary shifts in the median BGC-ome size across the phylogeny, we used an Ornstein-Uhlenbeck processes-based phylogenetic lasso approach via the *l1ou*^60^ R library (**Fig. 1c**). A set of 4,732 genus-representative genomes across the 74 orders were used to assess their functional profile and biosynthetic potential (**Supplementary Table S1**).

For additional details on the construction of the phylum-specific phylogenies, including construction with and without filtering for compositional heterogeneity, construction of the overall bacteria-archaea phylogeny, and details on parameter configurations for MCMCTree, please see the section with the same name in the **Supplementary Methods** document.

### Genome selection and annotation for systematic investigations of genetic traits and ecological trends correlated with bacterial biosynthetic potential

Species-resolution representative genomes were selected for each of the five bacterial phyla with the greatest number of characterized BGCs. Only species belonging to genera with either ≥ 5 (for Myxococcota), ≥ 10 (for Actinomycetota and Cyanobacteriota), or ≥ 20 (Bacillota and Pseudomonadota) genomes in GTDB R214 were considered (**Supplementary Table S3**)^61^. Different thresholds for different phyla were used to limit the number of genomes yet maintain an adequate sampling breadth for proper phylogenetic investigation of the focal phylum. For Bacillota, genomes belonging to related but distinct phyla in GTDB R214 (e.g. Bacillota_A which includes the genus of *Clostridioides*) were also included. A total of 17,844 species-representative genomes were assessed.

Functional annotation of coding genes for BGCs was performed using antiSMASH. Large scale annotations for known phylum-specific protein markers associated with multicellularity and sporulation^25–27,62–65^ were primarily performed using HMMER (v3.3.2)^66^ with profile HMMs gathered or constructed following *de novo* ortholog group inference across a more limited set of representative genomes^67^. SandPiper (v0.2.0)^68^ was used to extract ecological context based on metagenomic profiling for various GTDB taxa. For additional information on functional analysis of bacterial species-representative genomes, see the section with the same name in the **Supplementary Methods** document.

### In depth phylogenomic and evolutionary investigations of the phylum Actinomycetota

Ortholog groups were identified across a set of 133 genus-level representative genomes from the phylum of Actinomycetota using OrthoFinder (v2.5.4)^67^. The 133 representative genomes were a subset of those used for the phylum-specific phylogeny (**Fig. 2a; Supplementary Table S3-S4**) consisting of only assemblies which were designated “Complete” or “Chromosome” level for completeness status in NCBI’s Genome database. Coarse ortholog groups, rather than refined hierarchical ortholog groups, were used to prevent possible inappropriate paralogy splitting by OrthoFinder’s algorithm when applied to bacterial genomes (**Supplementary Table S6**).

A total of 34 ortholog groups were identified as being single-copy core (SCC), mostly corresponding to ribosomal proteins (26; 76.5%). Protein alignments for the 34 ortholog groups were generated using MUSCLE (v5.1), filtered using trimal (v1.4.1) in strict mode, and subsequently used as input for IQ-TREE for phylogenetic modeling of the phylum. IQ-TREE was run using an edge-unlinked partition analysis with ModelFinder limited to the LG and WAG substitution models and 1,000 bootstraps requested. The resulting phylogeny was largely topologically concordant with those based on a subset of the 74 universal bacterial genes by GToTree detection. The phylogeny was rooted by the genome for *Rubrobacter_D*, which was observed to be an early diverging genus in previously constructed phylum-specific phylogenies which included genomes from other phyla as outgroups.

Detailed methods for analyses pertaining to investigations of Actinomycetes evolution can be found in the document **Supplementary Methods**. This includes sections describing the D.stat metric for assessing topological models based on highly conserved sites of core proteins, the annotation of multicellularity and sporulation associated proteins, and comprehensive assessment of ansamycin BGCs and AHBA biosynthesis enzymes in 33,804 Actinomycetota genomes.

### Phylogenomic investigations of biosynthetic potential across the fungal kingdom

A total of 312 genomes were selected from across the fungal kingdom featuring representatives of different Dikaryon families, representatives of different classes from other fungal subkingdoms and phyla, or if they were genomes included in a recent study by Amses et al. 2022^35^, who sequenced several non-dikaryotic lineages of fungi, including many diverse chytrids (**Supplementary Table S7**). Representative genomes for Dikaryon families and non-dikaryotic classes were selected from species-level representative genomes which did not feature “fam_Incertae_sedis” or “unidentified” in their taxonomic designations in the Universal Fungal Core Genome (UFCG) database^69^ based on assembly N50. To properly assess biosynthetic potential, similar to our requirement for bacterial genomes, we required genomes to have a minimum N50 of 10kb. Four non-fungal eukaryotic genomes were included in analyses as outgroups: (GCF_000151315.2, *Capsaspora owczarzaki*; GCF_000001215.4, *Drosophila melanogaster*; GCA_000388065.2, *Fonticula alba*; GCA_000002865.1, *Monosiga brevicollis*). Genomes were downloaded using ncbi-genome-download and taxonomic names were systematically gathered from NCBI using taxonkit^70^.

Coding sequence (CDS) coordinates for assemblies were gathered in GFF format from NCBI’s RefSeq database or, if not available, from NCBI’s GenBank database. If CDS coordinate information was not available in either database or there were difficulties with using them to run antiSMASH for BGC predictions downstream (e.g. lack of CDS features or errors with overlapping CDS coordinates), then funannotate (v1.8.15; https://github.com/nextgenusfs/funannotate) was used to first mask repeat sequences, clean, and sort assemblies prior to performing gene prediction^71^.

UFCG profile (v1.0.5)^69^ was systematically applied to identify universal core protein sequences directly within genomic assemblies. Protein sequences for each of the 51 core universal gene markers found in ≥90% of genomes were extracted, aligned using MUSCLE, and trimmed in strict mode using trimal. Afterwards, these alignments were provided as input to IQ-TREE to construct a maximum-likelihood phylogeny using an edge-unlinked partition analysis with ModelFinder, allowing only for LG and WAG substitution models, and requesting 1,000 bootstraps. A monophyletic clade within Pezizomycotina enriched for BGCs was identified through visual examination and validated to correspond to classes described as enriched from prior studies^38,39^. For comparative analyses of biosynthetic genes or gene clusters across taxonomic groups, only a single representative genome per genus was regarded. Representative genomes were selected based on BGC-ome size.

For additional details on methods pertaining to further typing and assessing novelty of NRP synthetases or PK synthases^72,73^, annotation of HI associated domains and CAZymes, and the comparative and GWAS to understand enrichments genetic factors contributing to biosynthetic enrichment in Agaricomycetes and Pezizomycotina, please see the corresponding section in the **Supplementary Methods** document. **Visualization and select statistical tests**

Visualizations were primarily created using libraries in R and iTol^74^. Limits of boxplots shown alongside phylogenies correspond to 5th, 25th, 50th, 75th, and 95th percentiles, whereas stand-alone boxplots are always shown in Tukey’s style, where they extend to the largest value that is less than 1.5 times the interquartile range (IQR) from the top hinge of the box and the smallest value that is greater than 1.5 times the IQR from the bottom hinge of the box.

Spearman correlation coefficients and significance P-values of genetic and ecological quantitative measures with BGC-ome size were performed using the spearmanr function in scipy with a two-sided alternative model. One-sided rank sum tests were used to assess the statistical significance of differences in BGC-ome size between Actinomycetes with *ssgB*, Actinomycetes without *ssgB*, and other Actinomycetota without *ssgB*.

BGC-ome sizes and HI counts for genus-representative genomes were compared between different taxonomic groupings using one-sided Wilcoxon rank-sum tests. Taxonomic partitions within early-diverging fungal lineages were largely defined following taxonomic groupings highlighted by Amses et al. 2022^35^. Testing for enrichment of proteins with HI-associated domains within antiSMASH predicted BGC protocluster regions was performed using a one-sided Fisher’s exact test.

Additional details on other statistical models and tests can be found in the **Supplementary Methods**.

## Supplementary Notes

### Systematic investigations of the relation between bacterial lifestyle, genetic traits and ecological distributions with BGC-ome size

Of the 2,442 characterized BGCs cataloged in the MIBiG database^19^, 96.3% were from five bacterial phyla, Actinomycetota, Bacillota, Cyanobacteriota, Myxococcota and Pseudomonadota, and one fungal phylum, Ascomycota (**Fig. 1b**). To investigate why certain species encode for greater numbers of specialized metabolites, we assessed the relationship of BGC-ome size with other quantitative variables reflective of genomic, functional and ecological traits (**Extended Data Fig. 2ab; Supplementary Table S3**). Using a dataset of 17,844 species-representative genomes across the aforementioned five bacterial phyla, we reestablished that BGC-ome size was correlated with known factors such as genome size^75–77^.

The number of distinct carbohydrate utilization genes was broadly found to be the second most-associated factor with BGC-ome size. Genome-wide association studies performed individually for each phylum revealed individual CAZyme families associated with increased BGC-ome size^78^. These analyses revealed 75 families in Actinomycetota, 80 families in Bacillota, 56 families in Cyanobacteriota, 1 family in Myxococcota, and 92 families in Pseudomonadota as significantly associated with increased biosynthetic capacity. Only two families were found as significantly associated in four or more phyla, glycosyl hydrolase 19 and glycosyl hydrolase 19 subfamily 2, which correspond to a family of chitinases (**Extended Data Fig. 2c)**.

We also observed that genome-wide frequencies for some amino acids, as well as the number of distinct oxidative phosphorylation genes and adherence genes are consistently and significantly associated with larger BGC-omes in each of the five bacterial phyla. In contrast, some factors, such as the frequencies of other amino acids, were negatively correlated with BGC-ome size. In particular, the frequency of leucine was observed to be positively correlated with BGC-ome size, whereas the frequency of isoleucine was negatively associated. This trend might occur because the biosynthesis of leucine and coenzyme A, a key precursor of polyketides, share an enzymatic step that is not required for isoleucine synthesis^79^.

Genomic prediction of optimal growth temperatures further revealed that species with larger BGC-omes are mostly mesophilic (**Extended Data Fig. 2d**), which aligns with recent findings that mesophiles genomically encoded greater numbers of toxins^80^ and thermophilic fungi exhibit lower biosynthetic potential^81^. Further, of the bacterial genera which have at least ten characterized BGCs, many have large BGC-omes, often devote at least 300 kb in their genomes to specialized metabolism and are environmentally associated (**Extended Data Fig. 2e**).

### Evolutionary investigations to determine support for proposed phylogenetic model of Actinomycetes

In addition to our primary phylogeny for Actinomycetota based on single copy core ortholog groups, we constructed several alternate phylogenetic models. For some of the alternate models, multicellular Actinomycetes were discreetly placed within a single clade; however, further analyses suggest that their placement is likely a technical artifact due to them sharing greater numbers of genes with each other than with morphologically simpler genera (**Extended Data Fig. 4**).

Next, we developed a statistical framework to measure the number of conserved sites along protein alignments for single-copy core ortholog groups that supported morphologically simple families from clade-2 sharing a more recent last common ancestor with the *Streptomycetaceae* family than with families from clade-1 (**Fig. 2c**). This framework provided consensus support for our phylogenetic model in contrast to one where multicellular genera are aggregated within a single clade (**Fig. 2d; Extended Data Fig. 4**). Constructing a phylogeny of SsgB provided further support that the marker gene has largely evolved vertically, with instances from clade-1 and clade-2 genera mostly grouping separately (**Fig. 2d**).

We also searched for remnants of multicellularity-associated genetic markers, such as *ssgB* and *ramAB*^26,29,30,64,82,83^ across the entire Actinomycetota phylum. Several morphologically simple Acitnomycetia genera in clade-2 were found to harbor orthologs of multicellularity-associated markers, most notably chaplins, but not found outside of the Actinomycetes class (**Fig. 2a**).

Finally, a recent study identified a positive correlation between the extent of gene loss and mutation rates^84^. Ancestral state reconstruction analysis for ortholog group copy counts showed that genera with the greatest counts of cumulative gene loss were also phylogenetically the most distant from the last common ancestor of Actinomycetes along single-copy ortholog group-based core genome, providing yet additional support for the phylogenetic model we propose (**Fig. 2f**).

### Evolutionary analysis of anscamycin synthesizing BGCs and rifamycin resistance across Actinomycetota

Widespread rifamycin resistance has been predicted to occur across Actinomycetota and other phyla^33,85^. Some clinically used rifamycins are derived from a specialized metabolite produced by *Amycolatopsis* and *Micromonospora*, two highly divergent genera from clade-1 Actinomycetes. The metabolite belongs to the family of naphthalenic ansamycins, which includes a chemically homologous molecule, streptovaricin synthesized by some *Streptomyces* in clade-2. Homologous genomic regions to the rifmaycin encoding BGC from *Amycolatopsis* were identified across the Actinomycetota^86^ and used to establish a phylogeny for the cluster^87^ that was largely in alignment with our established species tree, thus largely in accordance with vertical evolution, despite high sequence conservation of core genes (**Extended Data Fig. 6a-d**). All homologous regions found belonged to the Actinomycetes class.

Similarly, the presence of the rifamycin-associated element (RAE)^85^ motif and homologs of the helicase HelR^33^, associated with rifamycin resistance were phylogenetically assessed and determined to be largely constrained to the class and found in both morphologically complex and simple genera from both clades (**Extended Data Fig. 6e-f**).

A major precursor of ansamycins is 3-amino-5-hydroxybenzoic acid (AHBA). One of the four key enzymes in the biosynthesis of AHBA is aminoDAHP synthase (RifH), which is related to DAHP synthases involved in the shikimate biosynthesis pathway. To better understand the evolution of aminoDAHP synthases, all DAHP synthases were extracted from Actinomycetota genomes and used to construct a phylogeny with RifH homologs from characterized ansamycin(-like) BGCs included^19^ (**Extended Data Fig. 6g**). The majority of inferred aminoDAHP synthases were found within a single clade of type II DAHP synthases. Further assessment of the taxonomic prevalence of different types of DAHP synthases and suspected RifH homologs in inferred AHBA gene clusters revealed that 90.5% of Actinomycetes genomes feature type II DAHP synthases (**Extended Data Fig. 6h**). In contrast, only 35.3% of other Actinomycetota genomes belonging to different classes featured type II DAHP synthases, which highlights the enzyme type as likely an important trait in Actinomceytia evolution. Identified RifH homologs were also exclusive to the Actinomycetes class, but rare, being found in <1% of Actinomycetes genomes.

## Supplementary Methods

### Construction of a bacterial and archaeal time tree and determination of evolutionary shifts in BGC counts

High-resolution phylogenies were independently constructed for the five bacterial phyla Actinomycetota, Bacillota, Cyanobacteriota, Myxococcota and Pseudomonadota using genus-resolution representative genomics. For each phylogeny, outgroup genomes from alternate phyla were included for rooting. The best hit for 74 proteins which are largely universal and found in single-copy across all bacteria were identified across genomes using GToTree (v1.7.07) in “best-hit” mode with otherwise default parameters ^51^. Some genera were dropped from investigation because they did not feature >50% of pre-computed single copy-core genes. Protein alignments were retained and filtered for sites where <10% of sequences featured gaps or ambiguous (“X”) characters. Full sequences which had gaps or ambiguous characters for ≥10% sites were also subsequently filtered. Retained protein alignments were then used as input to IQ-TREE (v2.2.0.3) to construct a phylogeny using an edge-unlinked partition analysis with models selected using ModelFinder limited to the LG and WAG substitution matrices and with 1000 bootstraps requested (https://github.com/Kalan-Lab/Ancestral_BGC_Exansions_Study/blob/main/Supplementary_Figures.pdf)^52,88^.

To assess the topological rigidity of the phylogenies constructed for the five phyla, we further filtered protein alignments to retain only 50% of the sites which are least impacted by compositional heterogeneity^89^. For each phylum, we first determined the median value of median pairwise GARP:FIMNKY ratios across protein alignments. This value was used as the GARP:FIMNKY ratio cutoff to discern between AT and GC rich genera. Afterwards, we computed, the degree of compositional bias per site as described in Muñoz-Gómez et al. 2019 and filtered for 50% of the sites with the lowest score. These filtered protein alignments were then used as input to IQ-TREE to construct a phylogeny once again using an edge-unlinked partition analysis. The topological structure of phylogenies achieved with and without accounting for compositional heterogeneity for each of the five phyla were largely congruent.

To construct a reliable and comprehensive phylogeny of bacteria and archaea, we first selected representative genomes for 77 bacterial and archaeal orders with greater than 200 genomes in GTDB R214 and which had a “Complete” or “Chromosome” quality assembly with an N50 > 100,000 from the major (most sequenced) genus to serve as a representative (**Supplementary Table S1**). The orders of Vampirovibrionales and Cyanobacteriales were automatically included to leverage valuable fossil information for construction of a timetree downstream. The orders UBA6257, which belongs to Candida Phyla Radiation^54,90^ and Fusobacteriales, which have been shown to be highly impacted by horizontal gene transfer^91^, were excluded to achieve clearer partitioning between bacteria and archaea and improve accuracy in phylogenomic modeling. GToTree was next used to identify the best-hit for a set of 16 universal ribosomal proteins^54^. The representative genome for the order of Nitrososphaerales was dropped for not featuring at least eight of the ribosomal proteins. The alignments for the ribosomal proteins were filtered for sites which featured gaps in fewer than 10% of genomes which resulted in retention of 1,095 amino acid sites along eight of the ribosomal proteins (L14, S10, S8, S6, L5, L2, S3 and S17). IQ-TREE was run using an edge-unlinked partition analysis and ModelFinder limited to the LG and WAG substitution matrices with a constraint tree manually constructed based on the phylum specific phylogenies constructed. Version v1.6.12 of IQ-TREE was used for this modeling instead of v2.2.0.3 to properly apply phylogenetic constraints. Targeted assessment of alternate phylogenetic placements of Myxococcales was additionally performed to resolve whether it was more appropriate to clade the order closer to Desulfobacterota or Pseudomonadota based on aggregate bootstrap support. The final phylogeny was rooted using the minimal ancestor deviation (MAD) technique^92^.

#### Bacterial and archaeal timetree construction

A concatenated alignment of the universal proteins was also created and formatted into Phylip format. CODEML (v4.10.0) was first used to estimate the Hessian matrix, gradient vector and maximum-likelihood estimates of branch lengths. The LG amino acid substitution model rate matrix, an alpha value of 0.5 for gamma rates and 5 discrete gamma categories were used to run CODEML. Afterwards, the *rst2* output file was renamed to *in.BV* and used as input for three replicate runs of MCMCTree to sample the posterior distribution for node ages. An approximate likelihood approach, an independent rate model and a uniform prior on phylogeny node ages were used to run MCMCTree. The parameters for the gamma-Dirichlet prior for the mean evolutionary rate were specified as =2, =40, =1, based on expectations from prior ancient phylogenetic investigations ^55,93^ and the parameters for the gamma-Dirichlet prior for the relaxed-clock model’s rate dispersion parameter were set to =1, =10, =1. The Monte Carlo Markov Chain (MCMC) was run with a burn-in of 100,000 and a sample size of 20,000 for 1,000 iterations. Results were assessed both visually and using effective sample size to confirm convergence in dating estimates^94,95^. All non-leaf nodes had an ESS of at least 4,141. TreeViewer^96^ and iToL^97^ were used to visualize results (**Fig. 1c; Extended Data Fig. 4**). For the iToL-based visualization in **Fig. 1c**, the standard deviation of the posterior age distribution was displayed as the width of branches.

#### Assessment of BGC-ome size differences across bacterial orders

To account for intra-order variation in BGC-ome sizes, genus-level representative genomes were selected for each of the 74 orders (**Supplementary Table S1**). These representative genomes were selected similar to a process described to select species-level representative genomes for the phylum specific analyses. The assembly with the greatest quality was selected based primarily on the assembly completeness category and secondarily on N50 as a measure of completeness, requiring representative assemblies to have a minimal N50 of 100,000. BGC and CAZy annotations of the genus-level representative genomes across the 74 orders were performed using the same approach we took for annotating species-level representative genomes for the systematic investigation of the five bacterial phyla (described above). An Ornstein-Uhlenbeck processes-based phylogenetic lasso approach, implemented as the function “estimate_shift_configuration” in the *l1ou*^60^ R library, was used to determine evolutionary shifts in the median BGC-ome size across the phylogeny (**Fig. 1c**).

Assessment of the number of distinct metabolites for each order in the phylogeny depicted in **Fig. 1c** were gathered from the Natural Products Atlas (using database files from 06/2023). The genus associated with each metabolite was mapped to orders based on the categorization of the genus in GTDB R214. Note, there is potential that some genera designations associated with metabolites might be inaccurate or have a different taxonomic classification if a genome was available for the source sample and classified using GTDB-Tk. The number of distinct metabolic nodes, which group metabolites more broadly compared to metabolic clusters, were calculated for each order.

### Genome selection and annotation for systematic investigations of genetic traits and ecological trends correlated with bacterial biosynthetic potential

To account for intra-genus variability in genomic content, representative genomes were selected for individual species (**Supplementary Table S3**). The process of representative genome selection for genera and species was primarily based on assembly completeness category and secondarily on assembly N50. Genus representative genomes were primarily used to establish phylogenies and no assembly quality filter was applied. However, an N50 ≥ 100,000 was required for species representative genomic assemblies to alleviate false negative detection of genetic traits^98,99^ and reduce partial detection of BGCs^100,101^. For some genera used for phylogenetic inference, no species representatives were selected. In certain instances, the most preferential genomic assembly selections for species or genera were no longer available for download on NCBI (largely due to authors of assemblies making them private retroactively). For these cases, the next best available genomic assembly was selected using iterative assessment. Assemblies were downloaded using ncbi-genome-download (v0.3.3)^102^ and gene-calling was performed using pyrodigal (v3.0.0)^103^ and prepTG (v1.3.9) to create GenBank files with CDS features^104^.

#### Functional annotations

BGC annotation was performed using antiSMASH (v7.0.0)^59^ with largely default settings, prodigal^105^ selected as the genefinding tool and GenBank files from prepTG processing as input. The BGC-ome size was computed as the summed length of all BGC regions identified whereas the NRPS+PKS-ome size was the sum of select BGC regions which feature NRPS or PKS associated types and at least one protocore region confidently predicted as a BGC (not necessarily NRPS or PKS). Phage and mobile genetic element coordinates were predicted using PhiSpy (v4.2.21)^106^ with the minimal number of phage genes reduced from 1 to 0 for detection. HMMs for carbohydrate utilization and transport genes from the dbCAN database (version 07262023)^107,108^ were downloaded and searched for in predicted proteomes using pyhmmer^109^ (v0.10.4). As recommended for annotation of bacterial genomes by the dbCAN curators, an E-value threshold of 1e-18 and a coverage greater than 0.35 were required for hits. Independent DIAMOND alignment (v2.0.15)^110^ searches in very-sensitive mode were used to identify homologs of transposons and insertion sequences from ISFinder^111^ (downloaded from https://github.com/thanhleviet/ISfinder-sequences on Dec 24th, 2023) and adherence associated proteins from VFDB (downloaded on Nov 17th, 2023)^112^. Prior to searches in target species-representative genomes, sequences for the two datasets were independently collapsed for redundancy using CD-HIT (v4.8.1)^113^ at 95% identity and 90% bi-directioal coverage (“-c 0.95 -aS 0.90 -aL 0.90”). Top hits from the databases were identified for proteins from representative genomes based on bitscore provided that alignments exhibited ≥30% identity and ≥70% bi-directional coverage. To account for functional redundancy, the number of distinct records for carbohydrate utilization and adherence genes identified within each genome was calculated. For instance, if multiple instances of a specific glycosyl hydrolase family were annotated in a genome (e.g. GH15), the count for the genome would only be incremented by one. Note, multiple dbCAN/CAZy profile HMMs could be represented by the same protein to accommodate for multi-domain protein architectures. The total count of proteins matching any CAZy profile per genome was also calculated. Insertion element counts were calculated per genome as the number of proteins shown to exhibit homology to sequences in the ISFinder derived database. Genes involved in oxidative phosphorylation were identified using KofamScan^114^ with KOfam HMMs associated with the KEGG pathway map00190. The unique count of KOfam HMM identified in each target genome was assessed to overcome complications with functional redundancy when correlating the measure with BGC-ome size. The predicted optimal growth temperature was calculated based on the proportion of specific amino acids using the formula described by Zeldovich et al.^115^.

#### Phylum-specific associations of CAZyme families with BGC-ome size

Pyseer (v1.3.11) was used to perform GWAS analysis to associate specific CAZyme families and sub-families with BGC-ome size. Specifically, a linear-mixed-model was applied, referencing pairwise similarities between genomes determined via phylogeny construction with GToTree^51^.

#### Phylum-specific annotations of markers associated with multicellularity and sporulation

An ortholog group from comparative genomics of representative Actinomycetota (see section: “In depth phylogenomic and evolutionary investigations of the Actinomycetota phylum”) was identified as corresponding to SsgB using a query protein from *Streptomyces venezuelae* (SCO1541). A profile HMM was constructed using HMMER (v3.3.2) and a reflexive search for the proteins used to construct the profile suggested that an appropriate score threshold for sensitive detection of the ortholog group was 44.5. This was used to identify additional instances of the ortholog group amongst all representative Actinomycetota genomes in GTDB R214. Social lifestyle-associated KOfam HMMs for Myxococcota were gathered from the study by Murphy et al. 2021^25^ and searched across all representative genomes from the phylum using KofamScan. Genes associated with sporulation formation in Myxococcota, previously determined using proteomics by Dahl et al. 2007^62^, as well as multicellularity and akinete formation in Cyanobacteriota, as described by Boden et al. 2023^27^ and Singh et al. 2020^63^, respectively, were searched using a strategy similar to what was described for identifying SsgB orthologs across Actinomycetota. For each phylum, OrthoFinder (v2.5.4) was run using default settings on a limited set of genus representative genomes. Profile HMMs were then constructed for coarse ortholog groups corresponding to proteins from the three studies and hmmsearch was used to comprehensively search all species-representative genomes. Score thresholds were determined by reflexive searches of profile HMMs on the sequences used to construct them.

*Assessment of ecological distribution*: The SandPiper (v0.2.0;

https://sandpiper.qut.edu.au/)^68^ database was assessed to determine the distribution of GTDB taxa across public metagenomes. SandPiper’s classification of metagenomes as either environmental or eukaryotic-host associated based on a gradient boosting approach was also used for specific analyses. For investigations of well-represented genera in the MIBiG database, if multiple partitions of a genus existed in GTDB, metagenomic presence for the canonical genus was assessed across all of the individual partitions in aggregate.

### In depth phylogenomic and evolutionary investigations of the Actinomycetota phylum

#### Annotation of BGCs, CAZymes and other functional traits

Distributions for genome size, distinct CAZy enzyme family counts and distinct oxidative phosphorylation enzyme counts were calculated for each genus based on data gathered for species-level representative genomes (**Supplementary Table S3)**. Distributions of the BGC-ome size were similarly based on antiSMASH annotations for species-level representative genomes. Consensus sequences were determined for orthologs using primarily MUSCLE in super5 mode (v5.1), or MAFFT with default options (v7.505)^116^ for a small subset of ortholog groups where MUSCLE experienced difficulties and HMMER (v3.3.2). Consensus sequences for ortholog groups were used to simplify functional annotation with EggNOG mapper (webserver v2.1.12) with *de novo* Pfam domain alignment requested (**Supplementary Table S5**)^117^.

#### Alternate phylogenomic and related investigations

To further assess the topology proposed in **Fig. 2ab** for the phylum of Actinomycetota, evolutionary model A, we applied alternate approaches for constructing a phylogeny of the phylum. First, we used IQ-TREE using ModelFinder, limited to substitution matrices WAG and LG, to generate individual phylogenies for the 34 single copy core genes which were then provided as inputs to construct a consensus network using SplitsTree4CE (v4.19.2)^118^ (**Extended Data Fig. 4a**). SplitsTree4CE consensus network construction was run using TreeSizeWeightedMean for edge weights, a threshold percent of 5.0 and a high-dimension filter applied. Rubrobracter_D was used to root the network. We also used a larger set of 84 near-SCC genes found in ≥ 80% of genomes and constructed a phylogeny similar to what was done for the strict-SCC set (**Extended Data Fig. 4d**). This phylogeny was in support of evolutionary model B, where Streptomycetaceae grouped with the majority of other mycelium forming Actinomycetes families from clade-1. To understand the discrepancy with our prior phylogenetic models, we further assessed the similarity in gene content between genomes. We developed *hamtree*, a program which accepts OrthoFinder results, computes the Hamming distance for shared ortholog groups between genomes and then uses these distances to construct a neighbor-joining tree. In the neighbor-joining tree constructed by hamtree, *Streptomycetaceae* and mycelium-forming clade-1 genomes were similarly grouped together in a monophyletic clade, suggesting the maximum likelihood phylogeny based on near-SCC ortholog groups might be affected by gene conservation in these taxa and large-scale gene loss in non-mycelium forming taxa (**Extended Data Fig. 4e**). Finally, we constructed protein alignments using MUSCLE for 186 core genes - ortholog groups found in all the genomes regardless of being single copy per genome or not. Two core ortholog groups were dropped because alignments could not be properly established without the removal of sequences. Next, individual ortholog group phylogenies were created once more using IQ-TREE using ModelFinder limited to substitution matrices LG and WAG which were then provided as input for STAG (downloaded on Sep 12, 2023)^119^ to determine a consensus species phylogeny (**Extended Data Fig. 4f**). The STAG based phylogeny was concordant with evolutionary model A for the phylum.

#### D.stat calculation

We developed the D.stat metric to assess genetic support between two phylogenetic models for the evolution of Actinomycetota based on a core protein alignment (**Fig. 2b-d**). The structure for this test was derived from Patterson’s D-statistic, used to detect signals of relatively recent introgression across a focal genome amongst related eukaryotic species ^120,121^. In our adaptation of the statistic, we similarly adopt a four leaf phylogenetic structure; however, we use a concatenated protein alignment instead of a whole-genome nucleotide alignment. The D.stat was computed for multiple combinations of four genomes which featured (i) one genome belonging to clade-2 that was not part of the family Streptomycetaceae, (ii) one genome from the family Streptomycetaea, (iii) one genome from clade-1 and (iv) an Acidimicrobiia genome. The use of a protein alignment is to account for the extreme evolutionary distances in the taxonomic relationships being investigated. Only biallelic sites along the concatenated protein alignment from 34 single-copy-core genes which featured the same residue in both Acidimicrobiia genomes were considered. Such conserved residues in the Acidimicrobiia outgroup were treated as ancestral alleles (*A*). The alternate allele was regarded as a derived allele (*B*). Under evolutionary model A, where clade-2 includes Streptomycetaceae as an early diverging sister of the remaining families, we would expect there to be a greater number of *AABA* and *BBAA* sites in comparison to *ABAA* and *BABA* sites (**Fig. 2bc**). As an example, an *AABA* site is one where the genome belonging to a family from clade-1 has a derived alternate allele, while the rest of the genomes - smaller genomes from clade-2 and the Streptomycetaceae genome - feature the ancestral allele observed in the outgroup Acidimicrobiia. In contrast, under evolutionary model B, where Streptomycetaceae groups with the majority of the clade-1 families capable of mycelium formation, we would expect the occurrences of *AABA*+*BBAA* and *ABAA*+*BABA* sites to be equal (**Extended Data Fig. 4bc**). Positive values for D.stat, supportive of the model A topology structure, were largely observed regardless of whether we used the single copy core protein alignment (feature 2,833 sites which are conserved amongst the outgroup Acidimicrobiia) or the near single copy core protein alignment (featuring 5,209 sites which are conserved amongst the outgroup Acidimicrobiia). Viewing D.stat distributions stratified by individual single-copy-core ortholog groups, revealed that some ortholog groups have distributions around 0 or have negative values, indicating either complex scenarios of horizontal transfer, the presence of convergent evolution, or Streptomycetaceae-specific differentiation from ancestral alleles common to the rest of Actinomycetes.

#### Detection of additional multicellularity-associated genetic markers and construction of ssgB phylogeny

The genomic carriage of multicellularity-associated genetic markers, SsgB^26,64,65^, SepX^83^, RamAB^29,30,82^ and ChpABCDEFGH^29,82^, are shown based on the presence of corresponding ortholog groups from OrthoFinder analysis (**Fig. 2a; Supplementary Table S5**). Literature reports of mycelium formation or complex clustering of cells was also manually gathered, primarily from Bergey’s Manual of Systematics of Archaea and Bacteria^28^ (**Supplementary Table S4**). A maximum-likelihood phylogeny for the ortholog group corresponding to SsgB was constructed using IQ-TREE with ModelFinder limited to substitution matrices LG and WAG and based on a protein alignment generated using MUSCLE in super5 mode which was trimmed using trimal in strict mode and filtered of a sequence which featured over ≥ 25% sites as gaps. AnGST (committed Aug 8th, 2016)^122^ was used to root the gene tree based on the species phylogeny (**Fig. 2e**).

*Relation between phylogenetic distance of single-copy core and gene-loss rates:* The count of ortholog groups at ancestral states was inferred using Count (downloaded from https://www.iro.umontreal.ca/~csuros/gene_content/count.html from Apr 3, 2010)^123^ with the asymmetric Wagner model. Because gene loss has been shown to be more common than duplications or gain in bacteria^124–126^, the gene gain-to-loss cost ratio parameter was set to two. Results from Count were parsed to determine the cumulative gene-loss count since the LCA of Actinomycetes for each Actinomycetes genome. The ete3 library^127^ was further used to determine the phylogenetic distance between the LCA of Actinomycetes and each leaf.

*Comprehensive assessment of the ecological distributions, resistance genes and characterized BGCs of representative families from Actinomycetota:* The 133 Actinomycetota genomes used for comparative genomics featured genera across 31 families. The ete3 library was used to collapse the SCC phylogeny of the phylum down to the family level (**Extended Data Fig. 5a**). To understand the habitat distribution of the 31 families, we assessed the number of metagenomes from each broad environment category they were found in at >0.01% relative abundance using SandPiper^68^. Metagenomes which corresponded to common metadata labels (e.g. groundwater metagenome) were manually binned into select categories (e.g. aquatic; non-marine). To determine the distribution of characterized BGCs from MIBiGv3.1^19^ across the Actinomycetota phylogeny, we extracted core biosynthesis proteins from their GenBank records and searched all 33,804 genomes belonging to the phylum in GTDB R214 using DIAMOND BLASTp with an E-value threshold of 1e-3 in fast mode. Characterized BGCs were regarded as present in a genome if >50% of core biosynthesis proteins were found, where each identified core biosynthesis protein exhibited a >95% identity and >70% coverage match. To sensitively assess antibiotic resistance traits across the phylum, which features many species for which resistance mechanisms might be currently poorly understood, we used the Core Resfams HMM set (v1.2)^128^ with HMMER and applied recommended gathering thresholds. Mean counts of each Resfam was calculated for each family across all genomes belonging to them.

#### Assessing the phylogenetic distribution and contexts of BGC associated ortholog groups

Key biosynthetic ortholog groups were identified if at least one member protein was found within an antiSMASH BGC region and marked as “rule-based-clusters’’ for the feature gene-function. Of these, 366 ortholog groups were identified as present within BGC contexts in genomes from both major clades of Actinomycetes and prioritized for further investigation. To more specifically associate ortholog groups with certain types of BGCs, the antiSMASH designated type of only the corresponding protoclusters it was found to overlap with were considered (**Fig. 2g**). This was important because BGC regions might include several independent protoclusters in BGC-rich taxa that are simply called as belonging to the same region due to genomic proximity^129^.

#### Assessing the evolutionary relationship between ansamycin encoding BGCs

A coarse and sensitive search for homologous BGCs to rifamycin (MIBiG entry BGC0000136) across all 33,804 Actinomycetota genomes from GTDB R214 was performed using fai (v1.3.11)^104^. fai was run requesting that homologous regions to the query BGC in target genomes possess matches to ≥50% of core proteins for biosynthesis with E-value thresholds of 1e-10 but loosened criteria for the percentage of total query genes. A subset of the homologous regions identified which featured ≥25% of the total genes from the query BGC at an average amino acid identity (AAI) of ≥60% were further examined through processing with zol (v1.3.11) to determine near single-copy core ortholog groups present in ≥90% of the gene clusters. Protein alignments for each ortholog group were produced using MUSCLE and trimmed for sites where ≥10% of sequences featured gaps using trimal. Ortholog group protein alignments were provided as input to IQ-TREE and run using the same settings that were used for constructing the species phylogeny.

Records for the characterized BGCs for synthesis of rifamycin from *Amycolatopsis* (BGC0000136), rifamycin from *Micromonospora* (BGC0000137) and streptovaricin from *Streptomyces* (BGC0001785) were obtained from MIBiG^19^ and investigated for homology and visualized using clinker^130^. Three genomes from the respective genera were further identified as possessing one of the three BGCs, based on high core gene similarity and ortholog groups between them were determined using OrthoFinder. OrthoFinder results were provided as input to paiof (v1.0.2) to determine the percent identity between core one-to-one ortholog BGC genes as well as all one-to-one orthologs between the genomes.

#### Assessing the distribution and conservation of rifamycin-resistance markers

The rifamycin associated element (RAE)^85^ was searched for across all 33,804 Actinomycetota genomes in GTDB R214 using BLASTn (v2.5.0)^131^ with the task mode set to blastn-short and restrictions set to perform only ungapped alignments with the following four 19-mers provided as queries: “GGGGCTTGCGGCAAGGCCC”, ”GGGCCTTGCGGCAAGGCCC”, “GGGGCTTGCGGCAAGCCCC”, “GGGCCTTGCGGCAAGCCCC”. Only exact hits which featured a sequence identity and coverage of 100% were considered (**Supplementary Table S6**).

The query sequence for HelR, a recently identified and widespread helicase contributing to rifamycin resistance^33^, was obtained from the CARD database (ARO:3007208; *Streptomyces venezuelae*)^132^. DIAMOND BLASTp was used to search for homologous proteins across the full set of Actinomycetota genomes from GTDB R214 in very-sensitive mode with an E-value threshold of 0.001. Alignments were loosely filtered for bi-directional coverage of ≥50% and sequence identity ≥30% to retain remote homologs and the corresponding target proteins were extracted. Only hits to proteins from genomes regarded as species-level representatives for the Actinomycetota phylum, previously selected for systematic investigations of variables associated with BGC-ome size, were considered. Multiple sequence alignment of the HelR homologous sequences was performed using MUSCLE super5 mode, trimmed using trimal in strict mode and used to create an approximate maximum likelihood tree with FastTree2 (v2.1.11)^87^.

#### Assessing the evolutionary origins of AHBA biosynthesis

The protein sequence for RifH, an aminoDAHP synthase, from the rifamycin BGC in *Amycolatopsis mediterranei* was used as a query to extract 10 homologs from 27 ansamycin(-like) BGCs from MIBiGv3.1^19^. 3-deoxy-D-arabinoheptulosonate 7-phosphate (DAHP) synthases were then identified across the full set of Actinomycetota genomes in GTDB R214 using KofamScan with the three KEGG ortholog (KO) profile HMMs corresponding to the enzyme. Classification of type I and II DAHP synthases^133^ was performed using hmmsearch with TIGRfam domains^134^ TIGR00034.1 (aroFGH; type I) and TIGR01358.1 (DAHP_synth_II; type II) as well the Pfam domain PF18152 (DAHP_snth_FXD), where the sequences were assigned to the best matching domain based on E-value. The aggregate set of DAHP synthase proteins was collapsed to remove redundancy using CD-HIT^113^ with parameters: “-c 0.80 -aS 0.90 -aL 0.90” and afterwards supplemented with the RifH homologs from MIBiG BGCs. Next, MUSCLE super5^135^ was used to align the remaining sequences, trimal^136^ was applied in strict mode to perform site-specific filtering and sequences with 125 or more gaps (threshold selected based on histogram of gap counts) were discarded prior to phylogeny construction of the remaining 2,923 sequences using FastTree 2^87^ and visualization using iTol^97^. fai^104^ was further used to identify co-located sets of proteins homologous to those involved in AHBA biosynthesis from the rifamycin BGC. A total of 1,539 gene clusters were regarded as involved in AHBA biosynthesis due to featuring three or more co-located AHBA biosynthesis proteins, RifGHIJKLMN, including at least two key proteins in the pathway (RifG, RifH, RifJ, RifK)^32^. Of these suspected AHBA biosynthesis gene clusters, all of which were found in Actinomycetes genomes, 582 featured an RifH homolog.

### Phylogenomic investigations of biosynthetic potential across the fungal kingdom

#### Timetree construction

A timetree was constructed by pruning the comprehensive phylogeny of 316 genomes for a subset of 212 fungal genomes featuring ≥55 UFCG universal gene markers per assembly in addition to the outgroup genome, *Capsaspora owczarzaki* (GCF_000151315.2) using ete3. The stringent requirement for the number of UFCG markers per genome was to construct a concatenated protein alignment to provide to MCMCTree. Across all 213 genomes, 22 UFCG were used to establish the strict core genome, where each gene contained sites that were ambiguous or gaps in at most 10% of genomes. The phylogeny was rooted by the *Capsaspora* outgroup genome and updated to encode priors for calibrations for MCMCTree analysis (**Supplementary Table S8**). MCMCTree analysis was performed using an identical methodology to what was previously described for constructing a timetree of bacterial and archaeal order representatives. To achieve convergence, the Monte Carlo Markov Chain (MCMC) was run with a burn-in of 20,000 and a sample size of 40,000 for 100 iterations in triplicate. All non-leaf nodes were confirmed to possess an effective sample size of at least 772 for the run used for dating estimates^94^.

#### Functional annotations

BGCs were predicted using antiSMASH^59^ with the taxon parameter set to fungi and GFF files describing CDS coordinates provided via the genefinding-gff3 argument. One predicted BGC region was discarded because codoff^101^ (v1.1.8) analysis and alignment of key biosynthetic proteins to bacterial RefSeq^137^ (downloaded on Nov. 4, 2021), suggested either extremely recent horizontal gene transfer from bacteria or contamination. BGC regions were regarded as (i) NRPS-like if they featured a protocluster categorized as “thioamide-NRP”, “NRPS”, “NRPS-like”, or “NRP-metallophore”, (ii) as PKS-like if they featured a protocluster categorized as “hgLE-KS”, “PKS-like”, “prodigiosin” “T1PKS”, “T2PKS”, “T3PKS”, “transAT-PKS”, or “transAT-PKS-like” and (iii) as terpene if they featured a protocluster categorized as “terpene”. The BGC-ome and strict NRPS-PKS-ome sizes of fungal genomes were computed similar to how they were for bacterial genomes. HMM profiles from the dbCAN database were searched for in predicted proteomes using pyhmmer. Hits were regarded if they featured an E-value of 1e-17 and a coverage greater than 0.45, as recommended for annotation of fungal genomes by the maintainers of dbCAN. The eggNOG-mapper server^117^ was used to perform comprehensive functional annotation of proteins belonging to select ortholog groups or hierarchical ortholog groups highlighted by the Pezizomycotina comparative genomic or GWAS analyses. Descriptions from eggNOG-mapper were manually assessed to infer functional categories. Homologous proteins which might function as heterokaryon-incompatibility proteins (HET) were identified using hmmsearch with gathering/trusted thresholds^66^ to search for the following Pfam profile HMMs: PF06985.15 (Heterokaryon incompatibility protein - HET; hit 9,067 CDS) and PF05729.16 (NACHT domain - NACHT; hit 5,270 CDS) (**Fig. 4d**).

#### Phylogeny construction and domain architecture assessments of HI-associated domains

PyHMMER was used to perform comprehensive annotation of Pfam domains to proteins with NACHT or HET domains. NACHT and HET domains from proteins were further extracted, independently aligned using MUSCLE super5^135^, trimmed in strict mode using trimal^136^ and used to infer approximate maximum-likelihood phylogenies using FastTree 2^87^. Domain architectures for HI-associated proteins were assessed based on comprehensive Pfam annotations using custom scripts provided in the associated code repository of this study.

*Orthology inference, comparative genomics of Pezizomycotina and biosynthetic potential GWAS in Dikarya*: OrthoFinder^67^ (v2.5.4) was used to infer ortholog groups between fungal genomes using gene annotations, as described above and default parameter settings.

Comparative genomics between Agaricomycetes and BGC-rich Pezizomycotina with the complementary set of all other dikaryon genomes was performed using coarse ortholog groups determined by OrthoFinder and a permutation-based approach. Briefly, 100,000 permutations with label shuffling were used to calculate empirical P-values in the mean copy count of ortholog groups between the two genome sets^138^. Only ortholog groups which were found in >70% of one group but <30% of the other group were investigated. All ortholog groups at such filters were highly significant, having the minimal possible empirical P-value. A total number of 124 ortholog groups were identified as significantly different in conservation or copy-count between the two groups (**Supplementary Table S9**).

The genome-wide association study (GWAS) to identify coarse ortholog groups associated with increased biosynthetic potential across the full set of 316 genomes was performed using a linear-mixed model in pyseer^78^ (v1.3.11). The BGC-ome size of genomes in Mbp units was treated as a continuous phenotype, an ortholog group presence or absence matrix was provided as the genotypes and the phylogeny constructed for the kingdom was pruned and converted into a kinship matrix using a script from the pyseer package to allow for population structure correction. After running pyseer, the count_patterns.py script within the pyseer package was used to determine an appropriate cutoff, accounting for multiple testings, for P-values adjusted for population structure. A total of 672 ortholog groups were identified as significantly associated with BGC-ome size (**Supplementary Table S10**).

Annotations for both ortholog group sets determined by the Dikaryon comparative genomics analysis and GWAS were performed using the eggNOG-mapper annotation server^117^. Protein sequences categorized as select ortholog groups were individually annotated with *de novo* searching for Pfam domains requested. The annotation with the largest score alignment score was selected as the best annotation for each ortholog group. Some additional annotations were performed for the set of 124 ortholog groups identified via comparative genomics using sequence alignment to the curated Swiss-Prot database^139^, via structural similarity using Phyre2^140^ and based on available literature.

#### Typing of PK synthases, NRP synthetases and key terpene biosynthesis genes

Protein sequences featuring either condensation domains from NRPS-like BGCs or ketoacyl-synthase domains from PKS-like BGCs were extracted and further typed using synthaser^72^ (v1.1.22) (**Fig. 3b**). Synthaser results were processed and used to determine the proportion of genomes for fungal taxonomic clades which featured different categories of type I PK synthases (non-reducing, partially reducing, or highly reducing), verify NRP synthetases and identify hybrid NRPS-PKS proteins. The proportion of genomes which featured suspected NRP synthases, possessing BGCs predicted as NRPS-like (antiSMASH types: NRPS, NRPS-like, NRP-metallophore and thioamide-NRP) with a condensation domain but not regarded as an NRPS by synthaser was individually determined as well. Similarly, the proportion of genomes which lacked a synthaser verified polyketide synthase but which featured a PKS-like BGC with a ketoacyl-synthase domain was also determined. NRPS-PKS and PKS-NRPS hybrid proteins were regarded as the same category. If BGC regions featured both NRPS-like and PKS-like regions, they were considered To similarly assess conservation of key terpene biosynthesis genes, we first identified six coarse ortholog groups from OrthoFinder analysis which were present in greater than 25% of fungal genomes and found as a key terpene biosynthesis protein in at least one genome by antiSMASH. After annotation of all proteins belonging to these six ortholog groups using eggNOG-mapper^117^, we identified five common Pfam domains across them: PF03936.20 (Terpene_synth_C), PF00348.21 (polyprenyl_synt), PF00494.23 (SQS_PSY), PF13243.10 (SQHop_cyclase_C) and PF13249.10 (SQHop_cyclase_N). Profile HMMs for these domains were gathered and systematically searched for in all proteins from across all genomes using PyHMMER^109^ with gathering cutoffs. For each terpene associated domain, the total proportion of genomes which feature the domain was reported regardless of whether hits were found within BGC contexts or not.

*Phylogenetic investigations of PKS ketoacyl-synthase domains and NRPS condensation domains from zoosporic and early-diverging fungi*: As described in the previous section, PK synthase and NRP synthetase sequences from NRPS-like and type I PKS-like BGCs featuring either condensation or ketoacyl synthase domains, respectively, were identified and processed through synthaser^72^. Synthaser results were processed and ketoacyl synthase domains were extracted from verified polyketide synthases (inclusive of hybrid NRPS-PKS proteins) belonging to genus-representative genomes from early-diverging lineages of fungi (fungi not belonging to Dikarya or Zygomycota). Similarly, condensation domains of non-ribosomal peptide synthetases were also extracted from early-diverging fungal genomes (also inclusive of hybrid NRPS-PKS proteins). The two sets of sequences were complemented with PKS ketosynthase domains and NRPS condensation domains from reference fungal BGCs cataloged in MIBiG (v3.1), which are primarily from Ascomycota and were similarly annotated for respective domains using synthaser. The protein sequences for the two categories of domains were then aligned using super5 mode in MUSCLE, trimmed with trimAl in strict mode and provided as input to FastTree 2 to infer phylogenies. Domain classes for sequences were determined using NaPDoS2^73^.

#### Phylogenetically standardized assessment of genome fluidity and pangenome size associated metrics

While genome fluidity is an informative metric which has been used to understand bacterial pangenomes^141–143^, it does not account for the core genome phylogenetic diversity represented within each taxonomic group making comparisons between different groups challenging. We thus developed psaps (https://github.com/raufs/psaps; v1.0.0) to assess informative metrics for pangenome expansion differences between taxonomic groups while accounting for phylogenetic diversity. Coarse ortholog groups determined by OrthoFinder and the core genome phylogeny of the fungal kingdom were provided as input to psaps. Briefly, by default, psaps calculates the summed branch lengths for subclades in the global phylogeny provided, by first pruning the global phylogeny to retain only leaves belonging to the focal subclade. It also computes the total number of distinct ortholog groups, auxiliary ortholog groups and the proportion of ortholog groups which are auxiliary for each subclade. The reason auxiliary ortholog groups are of interest is because core ortholog groups are likely to be present in the last common ancestor of the focal subclade and thus not informative for pangenome expansion assessment. The set of auxiliary ortholog groups per subclade is the total number of ortholog groups minus the core ortholog groups, where, by default, core ortholog groups are those found in at least 80% of genomes.

The psaps program also features an option to perform pairwise analyses between genomes independently for each subclade. This allows for the generation of a distribution of the various statistics for each subclade. In this mode the branch sum equates to simply the distance between pairs of genomes. If pairwise analysis is requested, the genomic fluidity metric^141^ and the standardized genomic fluidity metric are also calculated. The genomic fluidity metric for a pair of genomes belonging to the same subclade corresponds to the sum of the number of non-shared ortholog groups observed divided by the sum of the total number of ortholog groups observed in each genome. The genome fluidity is typically calculated for an entire microbial population or species by averaging values across multiple pairs of genomes; however, psaps reports values for individual pairs of genomes to allow for more resolute assessment of the distribution of values. The standardized genomic fluidity simply divides the genomic fluidity metric by the branch distance between the two genomes being assessed. This results in inflating the genomic fluidity metric when pairs of genomes are more similar to each other in the core genome phylogeny and deflating the metric when pairs of genomes are phylogenetically more diverged.

For the investigations of genome fluidity and BGC-ome size in **Fig. 4e** and **Extended Data Fig. 9**, a few genomes which had taxonomic classifications in NCBI that were discordant with expectations based on our or Amses et al. 2022 phylogenomic analyses were disregarded. Genomes removed for these analyses included: GCA_025526915.1 (*Entophlyctis*), GCA_025602905.1 (*Phlyctochytrium*), GCA_027604625.1 (*Blyttiomyces*). These genomes were still included in other analyses since their broader taxonomic classification (e.g. membership in the Chytridiomycota phylum) was reliable.

#### Aspergillus investigations of the relation between HI counts with BGC-ome and genome sizes

Representative genomes for 78 *Aspergillus* species with CDS predictions were downloaded from the NCBI Genomes database (**Supplementary Table S11**). They were annotated for BGCs using antiSMASH, CAZymes using PyHMMER with dbCAN profile HMMs and HI proteins using HMMER with HET and NACHT profile HMMs with analogous parameter configurations used for processing and analysis of the fungi-wide dataset of 316 genomes. A core genome phylogeny for the 78 species was next established using protein alignments of UFCG marker genes and OrthoFinder was applied to infer ortholog groups between the species, also using equivalent settings to what was performed for the fungi-wide dataset. Orthology predictions together with the core genome phylogeny and visually determined clade groupings for *Aspergillus* species, factoring in whether species exhibited similar counts of HI proteins, were provided as input for psaps to assess the relationship between the core genome phylogenetic breadth and number of auxiliary ortholog groups across different clades.

## Notes

### Competing Interest Statement

The authors have declared no competing interest.

