## Extended Data Fig. for "Complex multicellularity linked with expanded chemical arsenals in microbes"

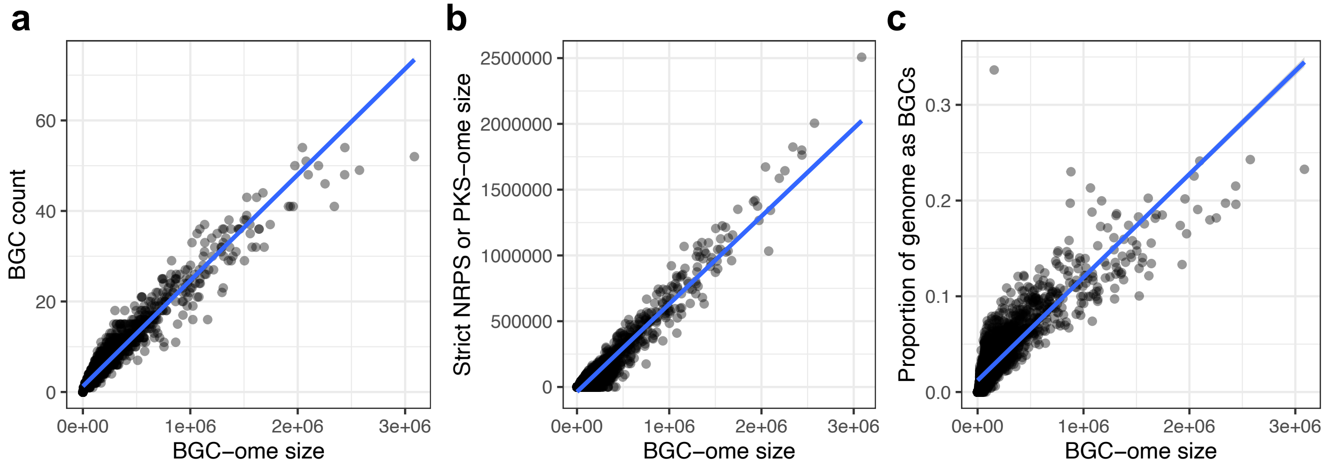


**Extended Data Fig. 1: Different metrics for assessing biosynthetic potential are highly correlated.** The relationships between BGC-ome size run using relaxed configurations in antiSMASH for 4,732 diverse bacteria and archaea with **a,** the count of BGC, **b,** the summed length of high-confidence BGC regions featuring NRPS or PKS protoclusters, and **c**, the proportion of entire genomes corresponding to BGCs.

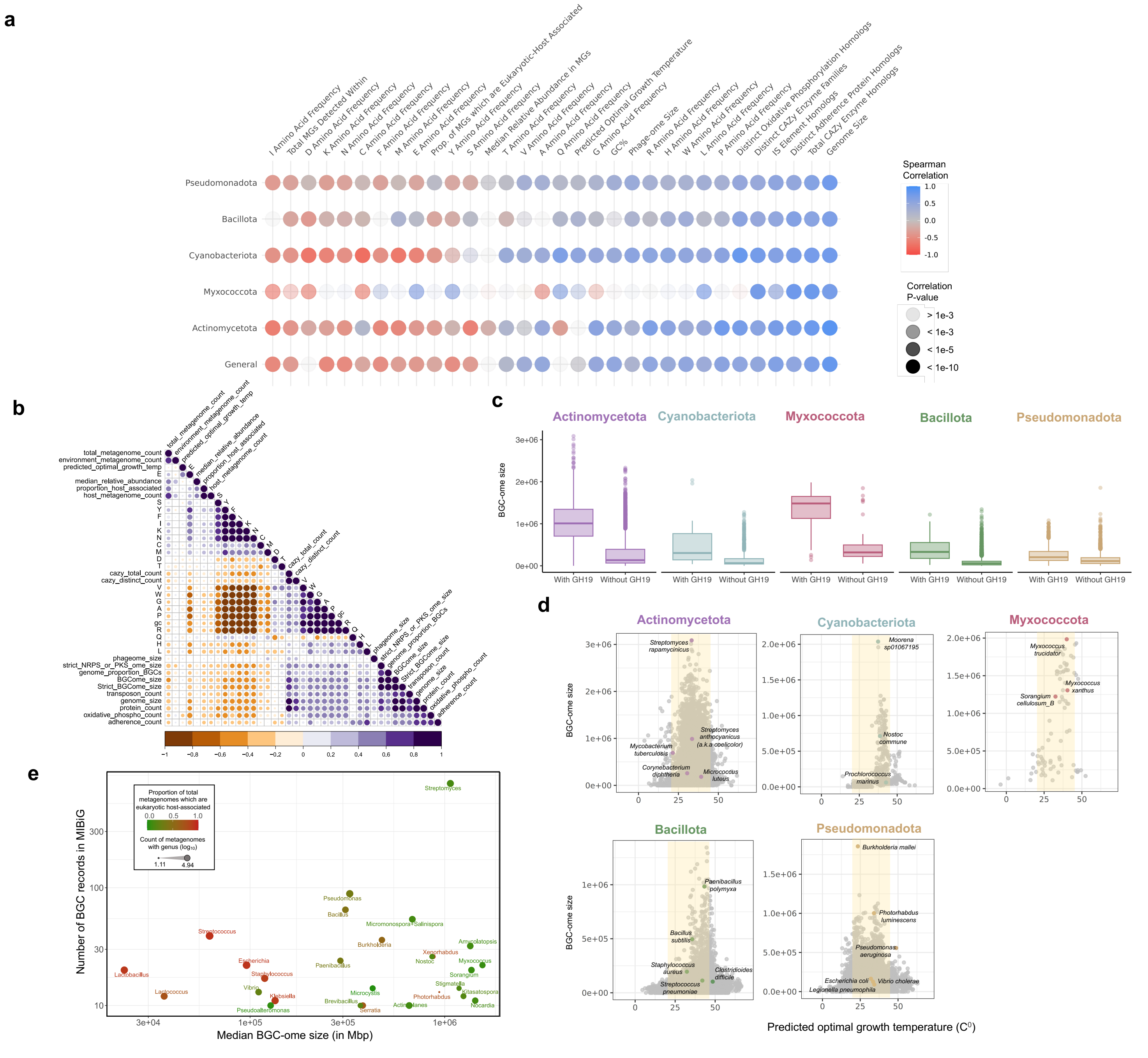


**Extended Data Fig. 2: Factors associated with increased biosynthetic capacity for the five bacteria phyla contributing most natural products. a,** Systematic relationships of various quantitative factors to BGC-ome size using Spearman's ranked correlation across all phyla (General) and for each of the five phyla individually. **b,** A lower-triangle heatmap showing the Spearman correlation between 38 quantitative variables. **c**, The BGC-ome size distributions of species-level representative genomes with and without glycosyl hydrolase family 19. For Myxococcota, the association was not significant after multiple testing correction. **d,** The relationship of BGC-ome size to genomic predictions for ideal temperature for growth of species in Celsius using the amino-acid frequency-based formula by Kurokawa et al. 2022. The yellow shaded area represents the temperature range for mesophiles (optimal growth temperature is between 20-45 C0). **e,** The relationship between median BGC-ome size and the number of MIBiG characterized BGC records for bacterial genera with ten or more characterized BGCs. Coloring corresponds to phyla, the size of the dots to the number of metagenomes the genus was inferred to be present in from the SandPiper database, and the shape corresponding to whether metagenomes genera were found in corresponded primarily to host-associated or environment, also based on the SandPiper database.

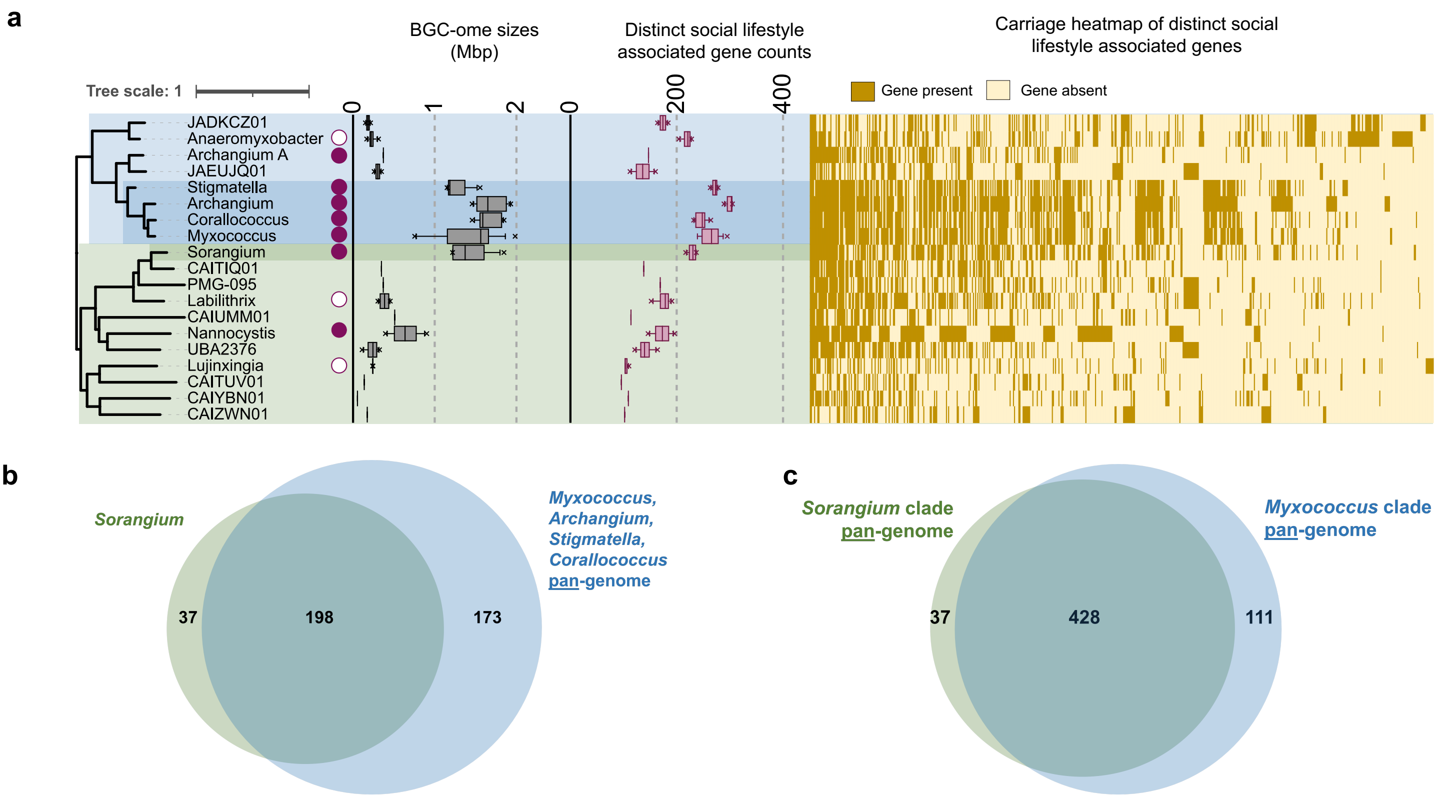


**Extended Data Fig. 3: Many genes contributing to sociality in Myxococcota are likely ancestral to the phylum. a,** A phylogeny of genus representatives from the Myxococcota phylum, with two major clades, is shown alongside whether corresponding genera have been experimentally validated as displaying a social lifestyle (fill circle = social lifestyle present, empty circle = social lifestyle absent; data from Murphy et al. 2021), distributions of their BGC-ome size and the count of distinct social lifestyle associated genes as described by Murphy et al. 2021 across species-level representative genomes. The presence or absence of 622 individual social lifestyle associated from the study as identified in the genus representative genomes is depicted as a heatmap. Venn diagrams show the overlap of distinct social lifestyle associated genes observed between **b,** *Sorangium* and at least one of four genera from a separate clade of the phylum, all of which are known to form fruiting bodies, as well as between **c,** their respective clades.

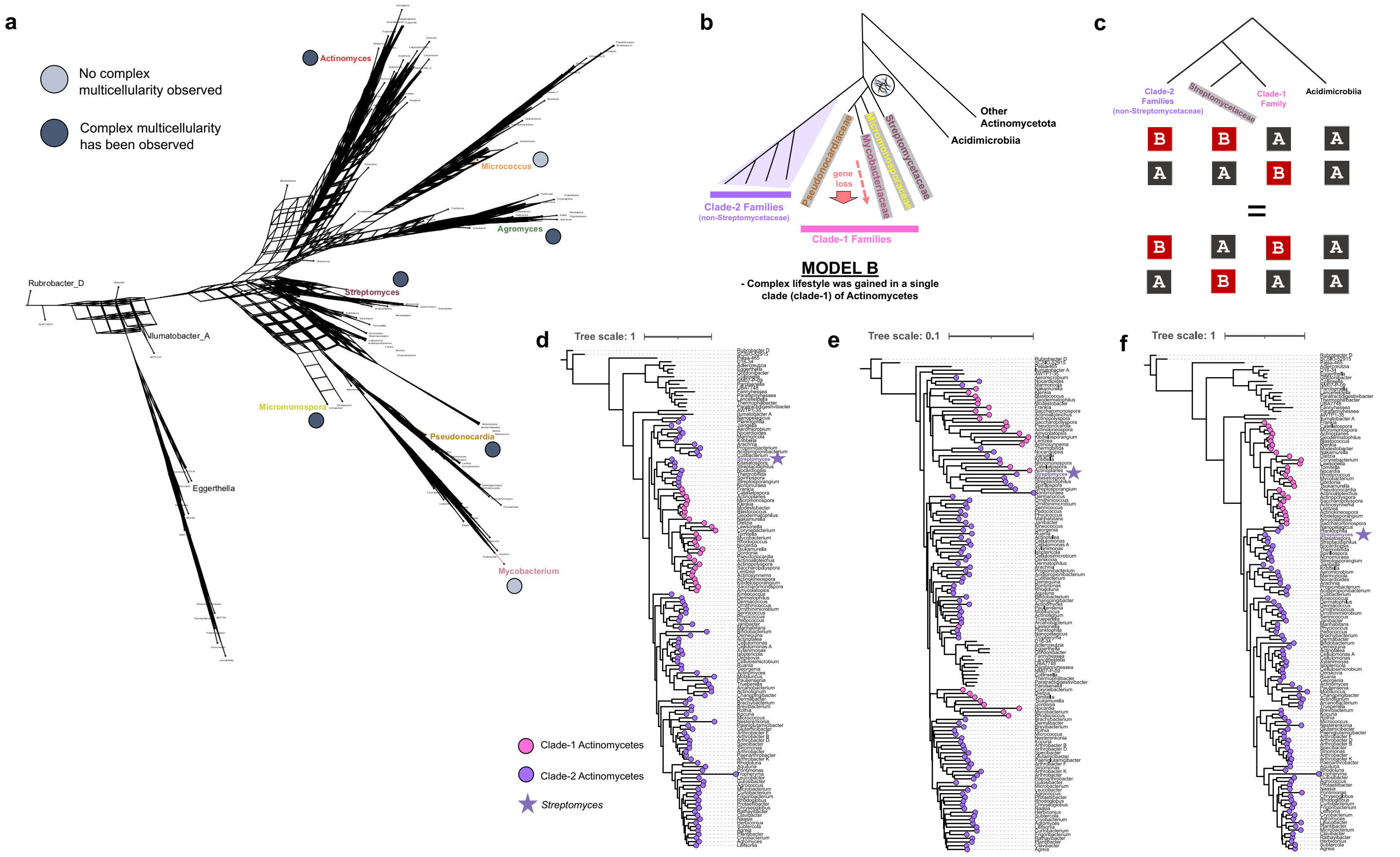


**Extended Data Fig. 4: Assessment of support for different topological models of Actinomycetota. a,** A phylogenetic network constructed from individual gene trees using SplitsTree4. Select genera are marked along with information on whether literature review indicates complex multicellularity exists for each such genus. **b,** An alternate topological model, Model B, for the evolution of the Actinomycetota phylum as proposed in Lewin et al. 2016 and by Gao and Gupta 2012. **c,** Under Model B, the ratio of BBAA & AABA sites to BABA and ABAA sites should be equivalent. **d,** A maximum-likelihood phylogeny constructed on alignments from 84 near (>80%) single-copy core is concordant with model B, placing *Streptomyces* into clade-1 with other morphologically complex Actinomycetes. **e,** A neighbor-joining tree based on Hamming distances of shared genes using hamtree also places *Streptomyces* with morphological complex Actinomycetes from clade-1. **f,** A consensus phylogeny based on individual gene trees for 186 core ortholog groups (include multi-copy genes) supports Model A.

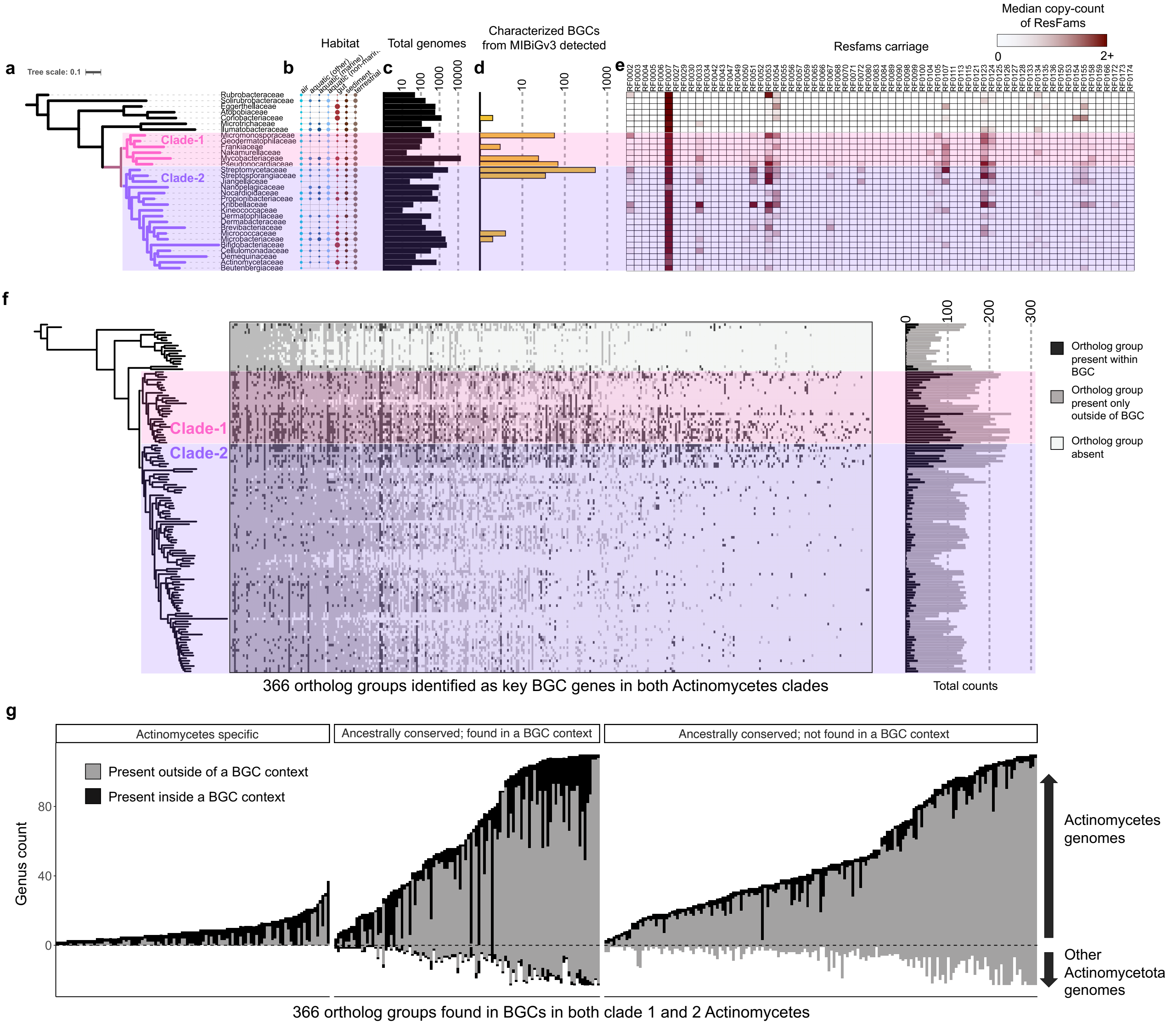


**Extended Data Fig. 5: Additional genetic, ecological, and biosynthetic information for Actinomycetota. a,** A collapsed version of the phylogeny presented in **Fig. 2a** where each leaf corresponds to a distinct family. **b,** Distribution of families across metagenomes from different habitats based on SandPiper derived data shown as a dotplot. **c,** The number of genomes within GTDB-R214 belonging to the family shown as a barplot (black). **d,** The number of distinct characterized BGCs from MIBiGv3 where at least 50% of key biosynthetic genes were detected at >95% amino acid identity being shown. **e,** The median copy-count for each Resfams resistance element is shown for families. **f**, Conservation of 366 ortholog groups found as core biosynthesis genes in both clade-1 and clade-2 Actinomycetes genomes is shown across a phylogeny of the Actinomycetota phylum. **g**, Details on the counts of genomes/genera with each of the 366 ortholog groups identified as corresponding to key protocore genes of BGCs in genomes from both major partitions of Actinomycetes. Upwards bars correspond to the number of Actinomycetes genomes with each ortholog group whereas downward bars indicate the number of non-Actinomycetes genomes with the ortholog group.

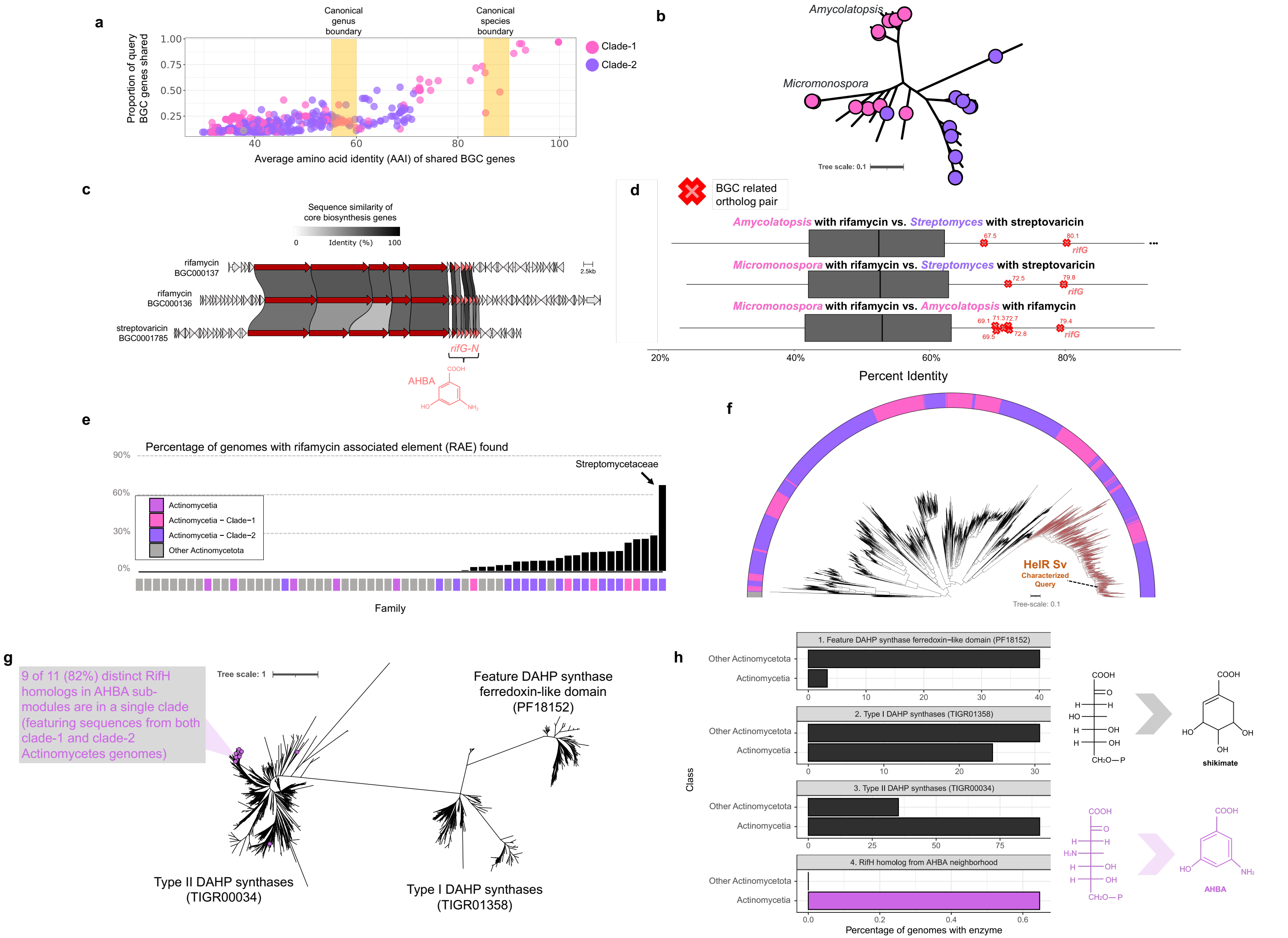


**Extended Data Fig. 6: Asamycin BGCs and rifamycin resistance elements are conserved across both major clades of Actinomycetes. a,** Average amino-acid identity plot from detected homologous regions to the rifamycin BGC [MIBiG entry BGC0000136] as a query against all Actinomycetota genomes in GTDB R214 using fai. **b,** A heterotachy-aware maximum likelihood phylogeny of near single copy core genes across homologous BGCs identified by fai to rifamycin exhibiting ≥ 60% AAI and ≥ 25% shared genes with the query BGC. **c,** Clinker-based illustration of sequence similarities of proteins from rifamycin BGC instances in divergent clade-1 Actinomycetes genera (*Amycolatopsis* & *Micromonospora*) and the streptovaricin BGC from *Streptomyces* in clade-2 Actinomycetes. **d,** Pairwise boxplots showcasing the range of amino-acid identities between orthologs using paiof for representative *Amycolatopsis*, *Micromonospora*, and *Streptomyces* genomes with either rifamycin or streptovaricin BGCs. The sequence identities for core biosynthesis genes from the BGCs are shown. Note, only one genome in GTDB R214 was identified to feature the streptovaricin BGC based on sequence similarity of core biosynthesis genes; however, the genome was of draft quality and the streptovaricin BGC was fragmented. Thus, only the identities of two core biosynthesis genes from streptovaricin to orthologs in rifamycin synthesizing BGCs in *Amycolatopsis* and *Micromonospora* are shown. **e,** The percentage of genomes within families across the Actinomycetota phylum carrying a rifamycin-associated element (RAE) sequence motif, often co-located with and involved in the regulation of rifamycin resistance enzymes, are shown. **f,** A comprehensive approximate maximum-likelihood phylogeny of homologous proteins of the rifamycin resistance contributing HelR enzyme from *Streptomyces venezuelae,* with the clade to which the query belongs to highlighted in red. The colored track indicates the taxonomic grouping of the genome from which the homolog was identified. **g,** A phylogeny of DAHP synthases from Actinomycetota with RifH homologs from MIBiG ansamycin(-like) BGCs marked in purple. **h,** Percentage of Actinomycetes or other Actinomycetota genomes with different types of DAHP synthases and inferred RifH homologs.

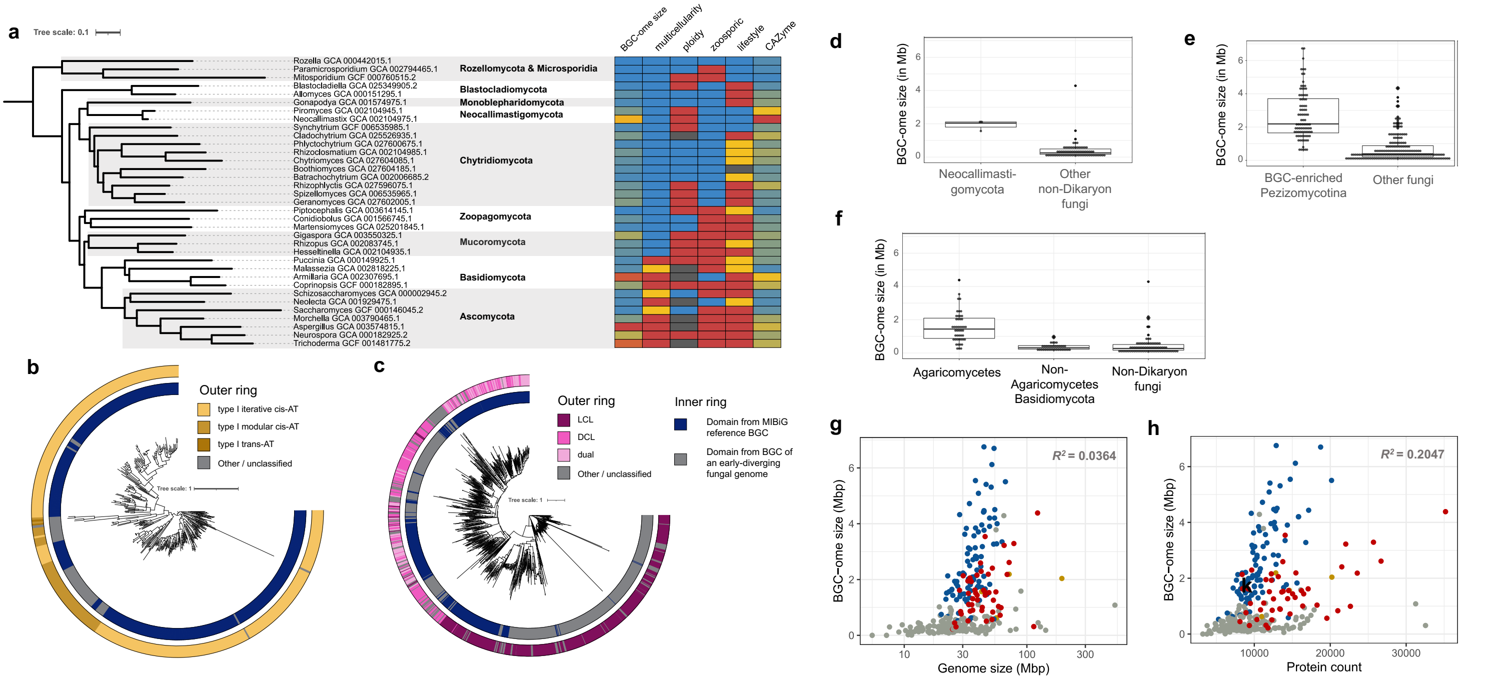


**Extended Data Fig. 7: Evolutionary and biosynthetic trends across the kingdom of fungi. a,** The phylogeny for the fungal kingdom pruned for 34 representative fungal taxa is shown alongside a heatmap depicting the species: BGC-ome size (from smallest to largest: blue [0], gold [1,811,838], red [3,623,676]), multicellularity type (complex multicellularity according to Nagy et al. 2018; yeasts (which can be dimorphic) = gold; simple multicellularity (e.g. sporangium / rhizoid formation) = blue), dominant ploidy status (haploid-dominant based on Amses et al. 2022 analysis or literature = red; diploid+ based on Amses et al. 2022 = blue; unknown = grey), zoosporic status (non-zoosporic = red; zoosporic = blue), lifestyle (free-living = red; host-associated / free-living = gold; endoparastic = blue), total CAZyme count (from smallest to largest: blue [0], gold [624.5], red [1,249]). Phylogenies for **b**, ketosynthase domains from PK synthases and **c**, condensation domains from NRP synthetases are shown with inner tracks depicting whether domains are from characterized BGCs in MIBiGv3.1 (dark blue) or early-diverging fungal genomes (non-Dikarya & non-Zygomycota; grey). Outer tracks show the domain class based on NaPDoS2 annotation. Comparisons of BGC-ome size are shown between **d,** Neocallimastigomycota and other non-Dikaryon fungi, **e,** BGC-enriched Pezizomycotina and other fungi, and **f,** Agaricomycetes, non-Agaricomycetes Basidiomycota, and non-Dikaryon fungi. The association of **g,** genome size and **h,** protein count with BGC-ome size. Genomes are colored in accordance with clade coloring in **Fig. 3a**. R2 values were computed using linear regressions adjusted for phylogenetic relationships between genomes.

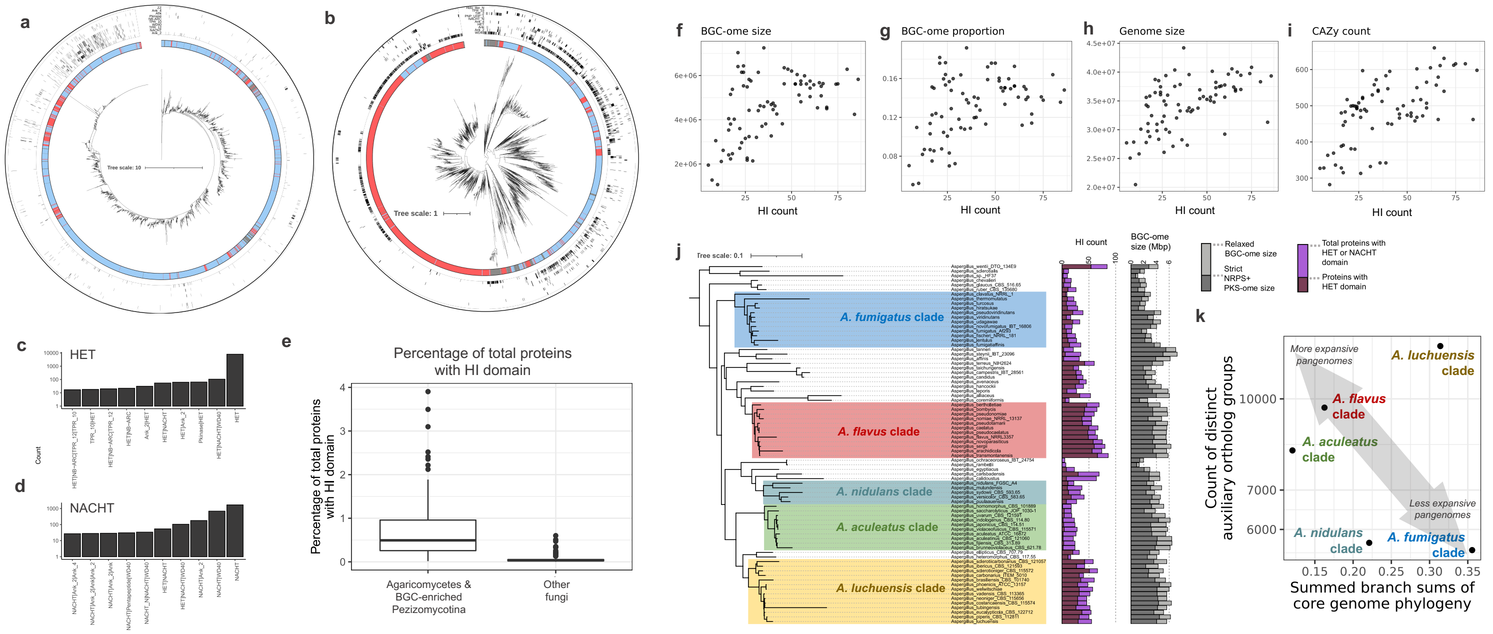


**Extended Data Fig. 8: Overview of HI proteins across fungi and associations with BGC-ome size and genome fluidity across the genus of *Aspergillus*. from fungal genomes.** Approximate maximum-likelihood phylogenies of **a,** HET and **b,** NACHT domains from proteins across the fungal kingdom. The inner-track indicates whether the domain is from BGC-enriched Pezizomycotina species (light blue), Agaricomycetes (salmon), or other fungi (grey). Co-occurrence information of the ten domains most commonly found co-occurring within proteins with the focal domain are shown as the outer-tracks. Frequencies for the ten most common domain architectures for proteins with **c,** HET and **d,** NACHT domains. **e**, The percentage of total proteins which are HI-related is shown for BGC-enriched clades in comparison to all other fungi is shown. The relationship between HI protein counts with **f**, BGC-ome size, **g,** the proportion of genome size corresponding to BGCs, **h,** the genome size, and **i**, the number of total CAZy homologs is shown across 78 genomes representative of different species belonging to the genus *Aspergillus*. **j,** The core-genome phylogenetic relationship of the 78 *Aspergillus* species is shown alongside barcharts depicting the total number of HI protein counts they have and their BGC-ome sizes. Clades were visually determined based on similar HI count distributions. **k,** The relationship between the number of distinct auxiliary ortholog groups (not belonging to the 80% loose core) with the summed branch length of clades from the core genome phylogeny is shown. Two clades with high HI counts, the *A. flavus* and *A. luchuensis* clades, had more expansive pangenomes than some clades with lower HI counts, such as the *A. fumigatus* clade.

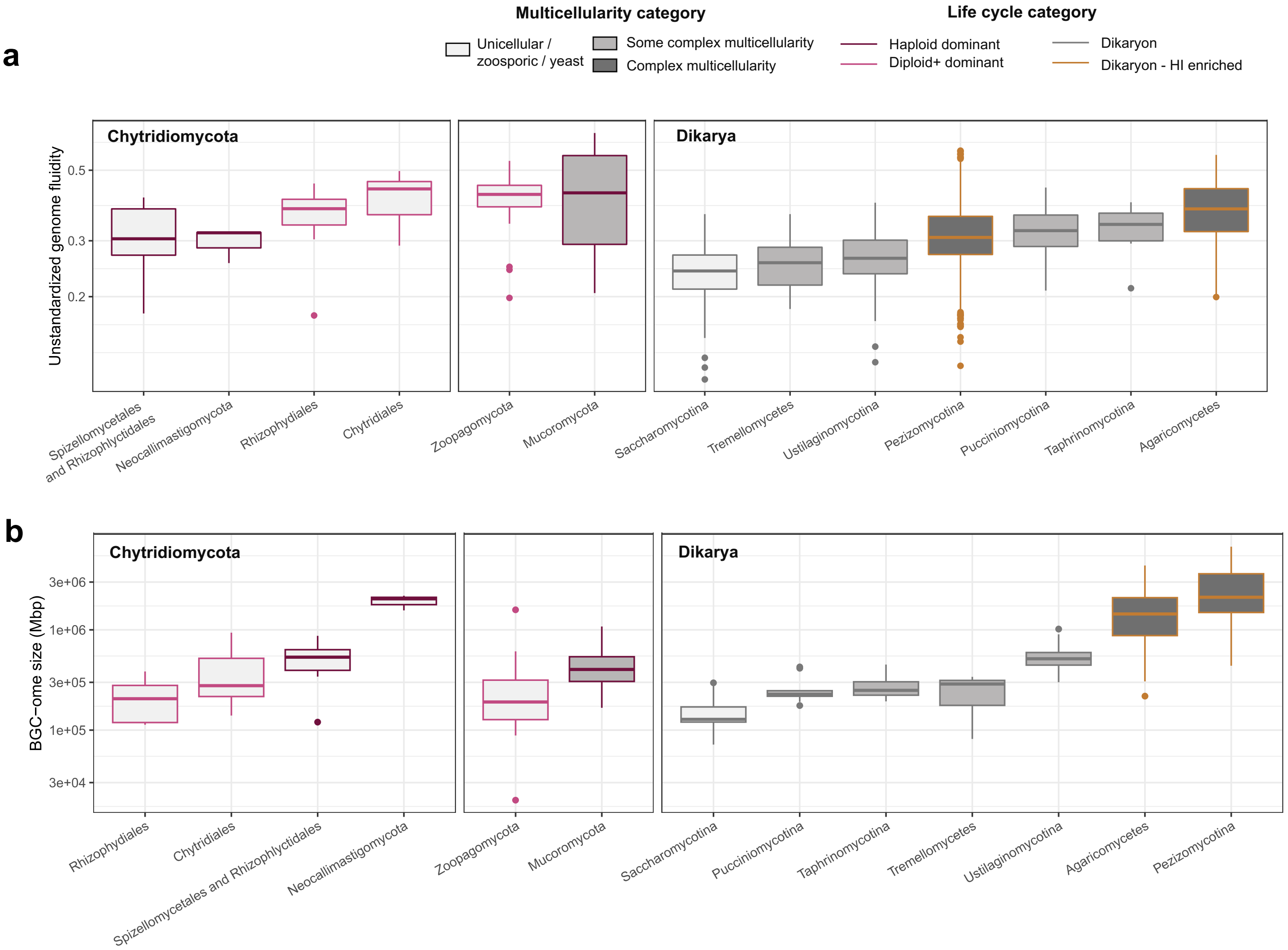


**Extended Data Fig. 9: Comparisons of unstandardized genome fluidity and BGC-ome size across fungal clades.** The distributions of **a,** unstandardized genome fluidity (not accounting for phylogenetic proximity between genomes) for pairs of genomes and **b,** the BGC-ome size per genome is shown for different fungal clades. Boxplots are colored and filled in accordance with the legend from **Fig. 4b.**
